# Transcriptional activation of *Bmal1* drives the inflammatory activity of monocytes by modulating mitochondrial unfolded protein response during hypobaric hypoxic stress

**DOI:** 10.1101/2024.04.02.587843

**Authors:** Yi-Ling Ge, Yong Liu, Bin Zhang, Jin Xu, Si-Yuan He, Qing-Lin Cao, Pei-Jie Li, Ying-Rui Bu, Yun-Gang Bai, Lin Zhang, Zhi-Bin Yu, Man-Jiang Xie

## Abstract

**Background:** Hypoxic stress-induced inflammation had been considered to play an important role in the onset and progression of altitude-related illnesses, but the origin of inflammatory cytokines, the specific responding cell types, and molecular mechanisms remain unknown. Mitochondria are responsible for oxygen consumption and recently reported to be the master regulators of inflammation, but it is not clear whether and how mitochondrial organelles sense the hypoxic stress and then control the inflammation.

**Methods:** Human subjects and mouse models were exposed to real or simulated altitude of 5500 m. Bone marrow-derived macrophages (BMDMs) and monocyte RAW264.7 cells were cultured under 1% oxygen hypoxic conditions. Myeloid-specific *Bmal1* knock-out mice were generated by crossing *Bmal1*^flox/flox^ mice with Lyz2-Cre mice. Inflammation was investigated by assessing inflammatory mediators, monocyte activities, and leukocyte infiltrating. Mitochondrial unfolded protein response was examined by measuring stress markers, such as LONP1, AFG3L2, and HSP60. The target molecular mechanisms were identified by performing bioinformatic analyses, ChIP assays, and gain/loss-of-function experiments.

**Results:** 1) Monocytes in peripheral blood mononuclear cell (PBMCs) were more sensitive and contributed promptly to circulating inflammation in response to acute hypobaric hypoxia. 2) Hypoxic stress triggered the mitochondrial unfolded protein response and then induced the mito-inflammation (NLRP3 inflammasome) in monocytes. 3) Activation of *Bmal1* drove mitochondrial stress and mito-inflammation by promoting Fis1-mediated mitochondrial fission in monocytes under hypoxia. 4) BHLHE40, a stress-responsive transcription factor directly targeted by HIF-1α, stimulated *Bmal1* transcription in monocytes under hypobaric hypoxia. 5) Myeloid-specific *Bmal1* deletion alleviated systemic circulating and vascular inflammation under acute hypobaric hypoxia.

**Conclusion:** BHLHE40, a transcription factor associated with hypoxia, stimulated *Bmal1*, which in turn triggered the mitochondrial unfolded protein response and drove the mito-inflammation in monocytes by promoting Fis1-mediated mitochondrial fission. Our work provides a novel mechanism which may develop the circadian targeting drugs for altitude or hypoxia-related diseases.

## Introduction

Altitude regions are of great significance in military, economic, recreational and religious activities^1^. With an increase in altitude, there is a decrease of oxygen pressure which results in the hypobaric hypoxic stress, posing a threat to individuals at high altitudes and causing high-altitude illnesses^2^. The most prevalent illness is acute mountain sickness (AMS) which could possibly progress to the more severe and potentially fatal conditions of high-altitude cerebral edema (HACE) and high-altitude pulmonary edema (HAPE)^3^. In addition, many common medical problems experienced by lowlanders, such as hemoglobinopathies, infections and chronic diseases of cardiovascular, gastrointestinal and pulmonary systems, are known to deteriorate during altitude travel^4, 5^. Recently, it has been demonstrated that hypoxia-induced inflammation in both systemic circulating and local tissue, particularly in the lungs and brain, plays a significant role in the progression of altitude-related illnesses^6–8, 9, 10^. However, the specific dynamic inflammation during short- and continuous-exposure to high-altitude and the specific cells responsible for producing inflammatory cytokines are not yet fully understood. Mononuclear phagocytes, a subgroup of immune effector cells, are bone marrow-derived myeloid cells which circulate in the blood as monocytes and migrate into tissues as macrophages during inflammation^11^. Monocytes are capable of producing inflammatory cytokines, engulfing cells and toxic molecules, and play an important role in mediating innate immune responses and inflammatory diseases^11, 12, 13^. The involvement of monocytes/macrophages in the inflammatory response under hypobaric hypoxia is rarely reported.

Mitochondria play a crucial role in controlling inflammatory response for containing multiple mitochondrial damage-associated molecular patterns (DAMPs) and activating some pattern recognition receptors (PRRs)^14^. The release of mitochondrial DAMPs could trigger inflammatory reactions by stimulating the cyclic GMP-AMP synthase (cGAS) and inflammasome signaling^14^, which serves as a precursor for numerous inflammatory diseases^15–17^. In addition, as the primary consumers of oxygen in cells, mitochondria could be significantly impacted by hypoxic stress, resulting in changes in mitochondrial dynamics and protein homeostasis accompanied with the mitochondrial dysfunction^18^. Mitochondrial unfolded protein response (UPR^mt^) is a mitochondrial specific stress response, in which stress indicators like mtDNA depletion or mutations, mitochondrial protein oxidation, or accumulation of unfolded proteins can trigger mitochondria-to-nuclear signaling to transcriptionally regulate the expression of mitochondrial matrix chaperones and proteases^19^. Mitochondrial stress has been reported to increase the levels of nod-like receptor 3 (NLRP3) inflammasome-dependent IL-1β and IL-18 in infected cystic fibrosis cells^20^. Therefore, it is postulated that mitochondria may transmit stress signals from hypoxia to mito-inflammation (an inflammation response related to mitochondria) by modulating the mitochondrial stress^14^.

Circadian clocks have been demonstrated to integrate the mitochondrial function and immune activities^24–27^. For instance, the core circadian gene *Bmal1* regulates monocyte movement by controlling mitochondrial homeostasis, leading to systemic inflammation and increased mortality in septic mice^28^. Circadian clocks convey temporal control to optimize the timing of fundamental cellular and biological processes following the environmental cues known as zeitgebers. After decades of studying, light, temperature, food, exercise and mechanosensory stimulation have been identified to be the zeitgebers. Recent studies have shown that fluctuations of oxygen levels can disrupted the circadian clock and there is an overlap (30%-50%) in synergistic activation between HIF1α and the core clock gene BMAL1^23^, indicating a fascinating interaction between hypoxic signaling and circadian rhythms^22, 23^.

The purpose of this study was (1) to investigate the dynamic inflammatory responses of circulating monocytes during short- and continuous-exposure to high altitudes in both human subjects and mouse models; (2) to determine whether hypoxic stress triggered the mitochondrial stress (UPR^mt^ and mitochondrial dysfunction) and the mito-inflammation (NLRP3 inflammasome activation) in monocytes; (3) to clarify the target molecular mechanism by which circadian gene *Bmal1* senses the hypoxia stress and subsequently triggers the mitochondrial unfolded protein response and mito-inflammation in monocytes.

## Methods

### Human subjects and study design

A total of 20 young (aged 19-25 years) male lowlanders, without history of smoking, cardiorespiratory disease, severe mountain sickness and recent exposure to altitudes >2000 m, were included in this before-and-after study. In general, human subjects were transported by trucks from the Kashgar region (1000 m above sea-level) to the peak of Karakoram Mountain (5500 m above sea-level) in stages. 5 ml peripheral venous blood was collected from each subject at 3 different altitudes and time points: (1) before departure at the altitude of 1000 m; (2) on the third day at the altitude of 5500 m; (3) on the 30^th^ day at the altitude of 5500 m. And in order to maintain the consistence of circadian rhythmicity, the blood collections at 3 different altitudes and time points all were conducted at 14:00.

The study protocol was approved by the Human Research Ethics Committee of Fourth Military Medical University and was conducted according to the Declaration of Helsinki principles. All participants gave their written informed consent to the study procedures after reading information about the study and having the procedures explained to them.

### Mice

Global Bmal1 knockout mice (Bmal1 ^−/−^ mice), Bmal1^flox/flox^ mice and Lyz2-Cre mice were generated on a C57BL/6 background and are commercially available from Cyagen Biosciences. Bmal1^flox/flox^ mice were crossed with Lyz2-Cre mice to yield mice with Bmal1 selectively deleted in myeloid cells throughout development and postnatal life. Wild-type mice (C57BL/6N) were purchased from animal experimental center of the Air Force Military Medical University (Xi’an, China). All mice were kept under 12 h light/12 h dark conditions with free access to food and water and the male mice at 8-12 weeks of age were used for experiments. Animal euthanization was completed using 3% pentobarbital for IP. All animal experiments were conducted according to the guidelines of the Institutional Animal Care and Use Committee of the Air Force Military Medical University.

### Cell Culture

RAW 264.7 cells were obtained commercially from the Pricella Life Technology (Wuhan, China, Cat#CL-0190) and cultured in Dulbecco’s Modified Eagle’s Medium supplemented with 10% fetal bovine serum (FBS) (Thermo Scientific, Rockford, IL, USA), 100 U/mL penicillin (Solarbio, Beijing, China), and 100 µg/mL streptomycin (Solarbio, Beijing, China). Bone marrow derived macrophage (BMDM) were isolated from mice tibias and femurs as described previously^29^ and cultured for 7 days in Iscove’s Modified Dulbecco’s Medium supplemented with 10% FCS (Thermo Scientific, Rockford, IL, USA), penicillin-streptomycin (100 U/mL and 100 μg/mL, respectively; Sigma-Aldrich) and 10 ng/mL murine macrophage colony-stimulating factor (M-CSF) (Peprotech).

For growth under hypoxic conditions, cells were grown in a specialized, humidified chamber (Heal Force, Shanghai, China) equilibrated with 1% oxygen / 94% nitrogen / 5% carbon dioxide for the indicated time.

### Collection of plasma and PBMCs

Human and mouse blood were collected and stored as described previously^47^. Approximately 5 ml human blood was collected with Ethylene Diamine Tetraacetic Acid (EDTA) as an anticoagulant for plasma extraction and then were stored at −80°C during transport until assay. Following venous blood collection, PBMCs were immediately isolated from peripheral venous blood by peripheral blood mononuclear cell isolation solution kit (Solarbio, Beijing, China, Cat#P6340) following the manufacturer’s instructions.

### Luminex liquid suspension chip detection

Luminex liquid suspension chip detection was performed by Cloud-Clone Corp. (Wuhan, China), and the Human Magnetic Luminex Assay kit (Cloud-Clone Corp., China) was used for in vitro quantitative measurement of factors in human plasma in accordance with the manufacturer’s instructions. All samples were run in a single plate per panel and in each sample of human plasma (n=20), with 3 immune mediators (IL6, IL-1β and CCL2) and 3 oxidative stress mediators (SOD, MDA and 8-OHdG) being analyzed.

### Detection of mitochondrial morphology and functions in cultured cells^30^

Mitochondrial morphology, ROS and mitochondrial membrane potential (MMP) in cultured cells (RAW264.6 cells and BMDM) were labeled by novel fluorogenic dyes of Mito Tracker ^TM^ Red CMROS (invitrogen by Thermo Fisher Scientific, Massachusetts USA, Cat#1800140), MitoSOX^TM^ Red mitochondrial superoxide indicator (invitrogen by Thermo Fisher Scientific, Cat#1842725) and Mito Tracker ^TM^ Red CMROS (invitrogen by Thermo Fisher Scientific, Cat#1800140) respectively, in accordance with the manufacturer’s instructions. Images were acquired using the Operetta CLS High-Content Analysis System (Harmony, PerkinElmer, Germany) with 63× Water/1.15 NA and the quantification of images fluorescence intensity was evaluated using PhenoLOGIC Harmony 4.9 supervised machine learning^51^.

### Detection of mitochondrial functions and Ly6C^Hi^ monocytes in PBMCs

Mitochondrial membrane potential (MMP), mitochondrial ROS and Ly6C in PBMCs were stained by fluorogenic dyes of Mito Tracker ^TM^ Red CMROS (invitrogen by Thermo Fisher Scientific, Massachusetts USA, Cat#1800140), MitoSOX^TM^ Red mitochondrial superoxide indicator (invitrogen by Thermo Fisher Scientific, Cat#1842725) and APC anti-mouse Ly6C antibody (Biolegend, Cat# 128015, 1:300 dilution) respectively, in accordance with the manufacturer’s instructions. Following staining, the sorting of monocytes in PBMCs and the detection of MMP, mitochondrial ROS, and Ly6C^Hi^ monocytes were performed by flow cytometric analysis (BD Bioscience, USA).

### Bioinformatic analysis

The gene expression profiles of the GSE13510, GSE19994, GSE148510 and GSE196728 datasets were downloaded from the NCBI Gene Expression Omnibus (GEO). Background correction, normalization, and expression calculation were performed using the robust multiarray average (RMA) method. Differentially expressed genes (DEGs) between groups were identified using empirical Bayes analysis with the Limma package, and a t-test followed by Benjamini and Hochberg (BH) adjustment was performed. Genes that met the cutoff criteria of a log (fold change) |log2FC|>1 and an adjusted p value < 0.05 were defined as DEGs. Gene Ontology (GO) term enrichment and Kyoto Encyclopedia of Genes and Genomes (KEGG) pathway analysis of the identified DEGs were performed using the databases of DAVID (Version 6.8) and Metascape (http://metascape.org/). Mitochondrial genes and mitochondrial pathways enrichment were identified by browsing MitoCarta3.0 (<u>MitoCarta3.0: An Inventory of Mammalian Mitochondrial Proteins and Pathways | Broad Institute</u>). The xCell algorithm was utilized in R software to analyze 64 immune cell types in human whole blood at high altitudes, employing and histograms and box plots. The proportions of immune cells in each group were evaluated via the Wilcoxon rank-sum test. A statistically significant difference was considered when p < 0.05.

### Lentiviral transduction

RAW264.7 cells were transduced with a lentivirus encoding *Bmal1* overexpression or control plasmid (SyngenTech, Beijing, China). The viruses diluted with complete medium to a final concentration of 1*10^8^ TU/mL before infection of RAW264.7 cells. Cells with stable overexpression of BMAL1 were screened with puromycin resistance and furtherly confirmed by western blotting analysis.

### Protein extraction and western blotting

Cells were dissociated by lysis reagent, and then protein was extracted. Protein was quantified using a BCA kit (Thermo Fisher Scientific, MA, USA) and western blotting was performed as previously described^31^. Briefly, equivalent amounts of proteins from different groups were loaded, electrophoresed and transferred to a polyvinylidene difluoride membrane (Millipore, Schwalbach, Germany). The membrane was blocked with 5% BSA in Tris-buffered saline with 0.1% Tween 20, followed by incubation with primary antibodies overnight at 4 °C. Then, the membrane was incubated with appropriate HRP-linked secondary antibodies for 1.5 h at room temperature. The membrane was finally imaged using an Odyssey scanner (LI-COR Biosciences, NE, USA) and analyzed using NIH ImageJ software. The antibodies used in this experiment are described in Supplementary Table S1.

### RNA Extraction and Real Time Quantitative Polymerase Chain Reaction (RT-qPCR)

Briefly, cells were homogenized with RNAiso (Takara, Otsu, Japan) and incubated in room temperature for 10 min. Then, the mixture was centrifuged at 12,000× g for 10 min at 4 °C, chloroform was then added to supernatants for phase separation. Total RNAs, located in the aqueous phase, were precipitated with isopropyl alcohol. After centrifugation, the supernatants were discarded, and RNA pellet was washed with 75% ethanol twice and dried for 10 min at room temperature. Finally, the pellet was dissolved in RNase-free water and stored at −80 °C for further analysis. Complementary DNA (cDNA) was amplificated with SYBR Premix Ex TaqTM (TaKaRa Bio, Otsu, Japan). RT-qPCR was performed using SYBR Green PCR master mix (Life Technologies). ACTB was used as the control for PCR product quantification and normalization. Data were analyzed via the relative Ct (2−ΔΔCt) method and were expressed as a fold change compared with the respective control. The sequences of primers for RT-qPCR are described in Supplementary Table S2.

### Ch-IP assay

Ch-IP assays were performed using the Ch-IP Assay kit (Thermo Fisher Scientific, MA, USA, catalog no. 26156). Cells were cross-linked with 1% formaldehyde for 10 min at room temperature and quenched in glycine. DNA was immunoprecipitated from the sonicated cell lysates using BMAL1 antibody and subjected to PCR to amplify the BMAL1 binding sites. The sequences of primers for Ch-IP assay are described in Supplementary Table S3.

### Plasmid and Oligonucleotides Transient Transfection

The DNA plasmid GV141-*Bmal1* (Genechem, shanghai, China), siRNAs targeting against *Bhlhe40* sequence (si-Bhlhe40), were used in the present study. The DNA plasmid and oligonucleotides transfection were performed using Lipofectamine3000 reagent (Invitrogen, Carlsbad, CA, USA) and Opti-MEM Reduced-Serum medium (Invitrogen) as described previously^31^. RAW264.7 cells were transfected with plasmid or oligonucleotides for 24 h before functional assays were carried out. The sequences of *Bhlhe40* siRNAs were 5’-AGAACGUGUCAGCACAATT-3’ (siRNA-1) and 5’-CCCUUCUCCUUUGGCACAUTT-3’ (siRNA-2).

### Multichannel fluorescence intravital microscopy (MFIM)

To observe the migration and infiltration of inflammatory cells into vasculature during hypobaric hypoxic stress, in vivo imaging of mice pulmonary vasculature combined with inflammatory cells (labeled by Alexa Fluor 488 anti-mouse CD11b antibody) was obtained through using the integrated imaging platform of MFIM (IVIM-MS model, IVIM Technology). WT and M-BKO mice were subjected to simulated high altitude of 5500m for 3 days by hypobaric chamber, and subsequently were injected with Evans Blue angiography agent and Alexa Fluor 488 anti-mouse CD11b antibody through the tail vein. After the angiography agent and antibody were diffused fully into the blood, the mice were anesthetized respectively, and the in vivo imaging was performed. During the imaging, 3 to 4 mice were used in each group, and the pulmonary vasculature of each mouse was observed at multiple time points, with representative images and videos being taken.

### Statistical Analysis

For statistical analysis, all quantitative data are presented as the mean±SEM. Statistical analysis for comparison of 2 groups was performed using 2-tailed unpaired/paired Student t tests. Statistical differences among groups were analyzed by 1-way ANOVA or 2-way ANOVA (if there were 2 factor levels), followed by Bonferroni’s post hoc test to determine group differences in the study parameters. All statistical analyses were performed with Prism software (GraphPad prism for Windows, version 9.0, Nashville, TN). Differences were considered significant at *P<0.05, **P<0.01, and ***P<0.001^32^.

## Results

### 1. Hypobaric hypoxia induced a dynamic inflammatory response of plasma and PBMCs from human subjects and animal models during 3-day and 30-day exposure

In order to investigate the dynamic inflammatory response, a human study was conducted with 20 male volunteers. These subjects were transported by truck to climb the Karakoram Mountain (altitude of 5500 m), and their peripheral blood samples (5 ml/person) were collected at different time points. As the baseline reference, the first collection was established before departure at the altitude of 1000 m. The second collection was done on the third day at an altitude of 5500 m in Karakoram Mountain, followed by collections on the 30^th^ day (Fig. 1A). The circulating inflammatory cytokines in plasma were measured using the Luminex assay. As compared with that of the baseline of 1000 m, the plasma levels of IL-1β, MCP-1 and IL-6 significantly increased on third day at the altitude of 5500 m and continuously increased on 30^th^ day at the altitude of 5500 m (Figure. 1B), suggesting that acute and continuous exposure to high-altitude hypoxia triggers an obvious circulating inflammatory response.

**Figure 1.**
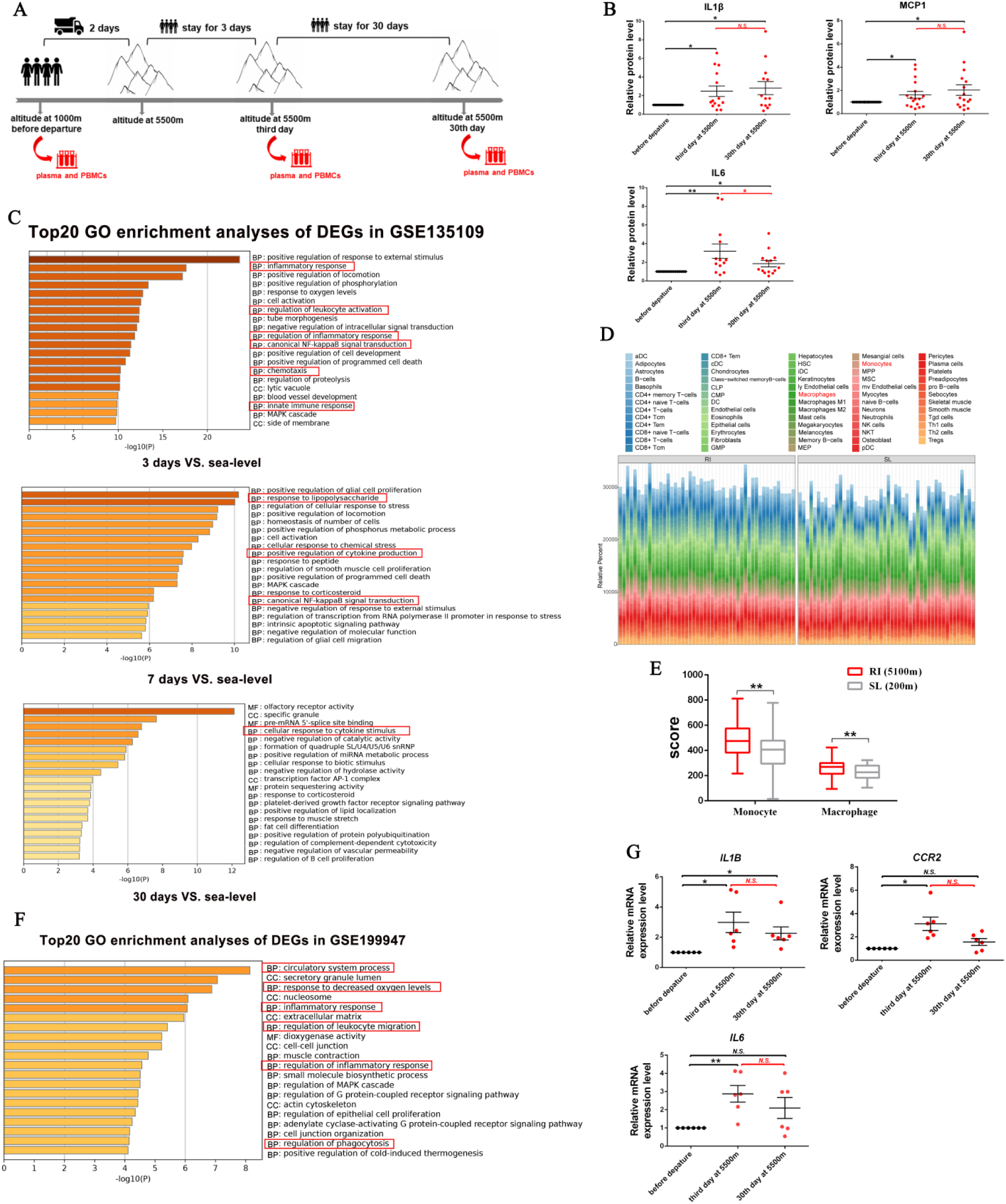
The dynamic inflammatory response of plasma and PBMCs from human subjects in 3-day and 30-day exposure to hypobaric hypoxia. **A**, Scheme of human experiments: human subjects were recruited to climb the Karakoram Mountain by truck, with their plasma and peripheral blood mononuclear cells (PBMCs) being collected at three different time points: on the third day at the Karakoram Mountain peak (∼5500m); on the 30th day at the Karakoram Mountain peak (∼5500m); and, as a baseline reference, before departure at the Kashgar region (∼1000m). **B**, Dynamic expression levels of inflammatory cytokines (IL6, IL-1β, and MCP-1) in human plasma during the process of altitude climbing, by Luminex assay (n=20; normalized for nonspecific binding). **C**, Top20 GO enrichment analyses of DEGs in the human peripheral leukocytes between sea-level group and three high-altitude groups: staying at the Qinghai-Tibet Plateau for 3, 7 and 30 days, respectively. Red frames indicate the inflammation related pathways (n=3; source data were obtained from GSE135109). **D**, Landscape of the abundance of 64 immune cell types in human whole blood at La Rinconada (RI, Peru 5,100m) (n=8; source data were obtained from GSE196728). **E**, Box plots of the monocytes and macrophages proportions in human whole blood at RI, relative to at sea-level (200m) (n=8; source data were obtained from GSE196728). **F**, Top20 GO enrichment analyses of DEGs in human monocytic THP-1 cells between normoxia group (n=3) and hypoxia group (under 1% oxygen hypoxic condition for 8 and 72h; n=6). Red frames indicate inflammation related pathways. (source data were obtained from GSE199947). **G**, Dynamic mRNA expression levels of inflammatory cytokines (*IL6*, *IL-1β*, and *CCR2*) in human PBMCs during the process of altitude climbing, by RT-qPCR analysis (n=8). **B and G**, Data represented as mean± SEM. **E**, in boxplots, center indicates the median, and the bottom and top edges indicate the 25th and 75th percentiles, respectively. *P<0.05, **P<0.01, *** P<0.001, **** P<0.0001, *N.S.* no significance. Statistical significance was determined by paired Student’s 2-tailed t test (B and G), and the Wilcoxon rank-sum test (E).

Inflammatory cytokines are predominantly released from leukocyte immune cells, including monocytes, macrophages, and lymphocytes. Based on the GSE135109, the bioinformatic analysis was used to identify the role of inflammatory pathways in the leukocytes at the high altitude of Qinghai-Tibet Plateau for 3, 7 and 30 days, respectively (Fig. 1C and Fig. S1A), which identified 5629, 4787 and 9824 differential expression genes (DEGs) in the three high-altitude groups, respectively. Gene Ontology (GO) enrichment analysis indicated that 3-day acute and 7-day continuous exposure to high-altitude significantly induced the leukocyte activation, cytokine production, and transduction of NF-κB signaling (Fig. 1C, red frames); while prolonged exposure to 30 days obviously reduced the inflammatory pathways in human leukocytes (Fig. 1C, red frames). Next, in order to identify the specific source of circulating cytokines in response to high-altitude hypoxia, xCell analysis based on GSE196728 was performed to investigate the abundance of 64 types of human immune cell in the regions of La Rinconada (altitude of 5100 m) (Fig. 1D); and it was confirmed that the proportions of monocytes and macrophages were significantly greater in the immune cell types at high altitude (Fig. 1E and Fig. S2). In addition, GO enrichment analysis of GSE199947 indicated that in human monocytic THP-1 cells, a number of inflammatory genes were significantly increased in response to decreased oxygen levels (Fig. S1B), which suggested an important role of monocytes in the hypoxia-induced inflammation (Fig. 1F, red frames). In the present work, the monocytes-enriched peripheral blood mononuclear cells (PBMCs) were isolated from the human subjects to assess the expressions of inflammatory cytokines by RT-qPCR analysis. As shown in Fig.1G, the mRNA expression levels of IL-1β, CCR2 (the receptor for MCP-1), and IL6 in PBMCs significantly increased on the third day at 5500 m; IL-1β expression continuously increased on the 30^th^ day at 5500 m, but CCR2 and IL6 expression decreased into original level on the 30^th^ day at 5500m. Together, our results clearly indicated that the levels of circulating cytokines increased in 3-day acute exposure to high altitude and maintained at a high level in the prolonged exposure to 30 days. In addition, the human monocytes-enriched PBMCs showed a significant inflammatory response in acute exposure to high altitude, which may be responsible for the dynamic changes of circulating cytokine when acute and prolonged exposure of hypobaric hypoxia

### 2. Hypobaric hypoxia induced the dynamic mitochondrial stress and mito-inflammation in PBMCs from human subjects and mouse model studies during 3-day and 30-day exposure

To screen the inflammatory pathways of monocytes in response to hypobaric hypoxia, Cellular Component enrichment analysis based on GSE135109 indicated that a large proportion of DEGs were located in mitochondria when exposed to high-altitude for 3 days and 7 days (Fig. 2A, red frames). Furthermore, mitocarta3.0 database, an inventory of mammalian mitochondrial proteins and pathways, suggested that exposure to high altitude obviously induced 118 mitochondrial DEGs (Fig. S1C), which were closely related to the mitochondrial protein homeostasis (Fig. 2B, red frames). Therefore, the present work was to investigate the mitochondrial oxidative stress markers (SOD, MDA, and 8-OHdG), mitochondrial unfolded protein response (LONP1, AFG3L2, and HSP60), and mito-inflammation (NLRP3) in the plasma and PBMCs of human subjects (Fig. 2C and Fig. 2D). As indicated by the Luminex assay, the plasma levels of SOD and 8-OHdG began to significantly increase on 3-day exposure of 5500 m and still maintained relatively high levels on 30-day exposure of 5500 m; the plasma levels of MDA began to rise on 3-day exposure of 5500 m and reduced to the baseline level of 1000 m on 30-day exposure of 5500 m (Fig. 2C). RT-qPCR analysis indicated that both the markers of mitochondrial unfolded protein (LONP1, AFG3L2, and HSP60, Fig. 2D) and NLRP3 (mito-inflammation, Fig. 2E) in PBMCs significantly increased on 3-day exposure of 5500 m and then decreased to the baseline level of 1000 m on the 30-day exposure of 5500 m, which indicated that mitochondrial stress and mito-inflammation are more sensitive in response to acute exposure of high altitude.

**Figure 2.**
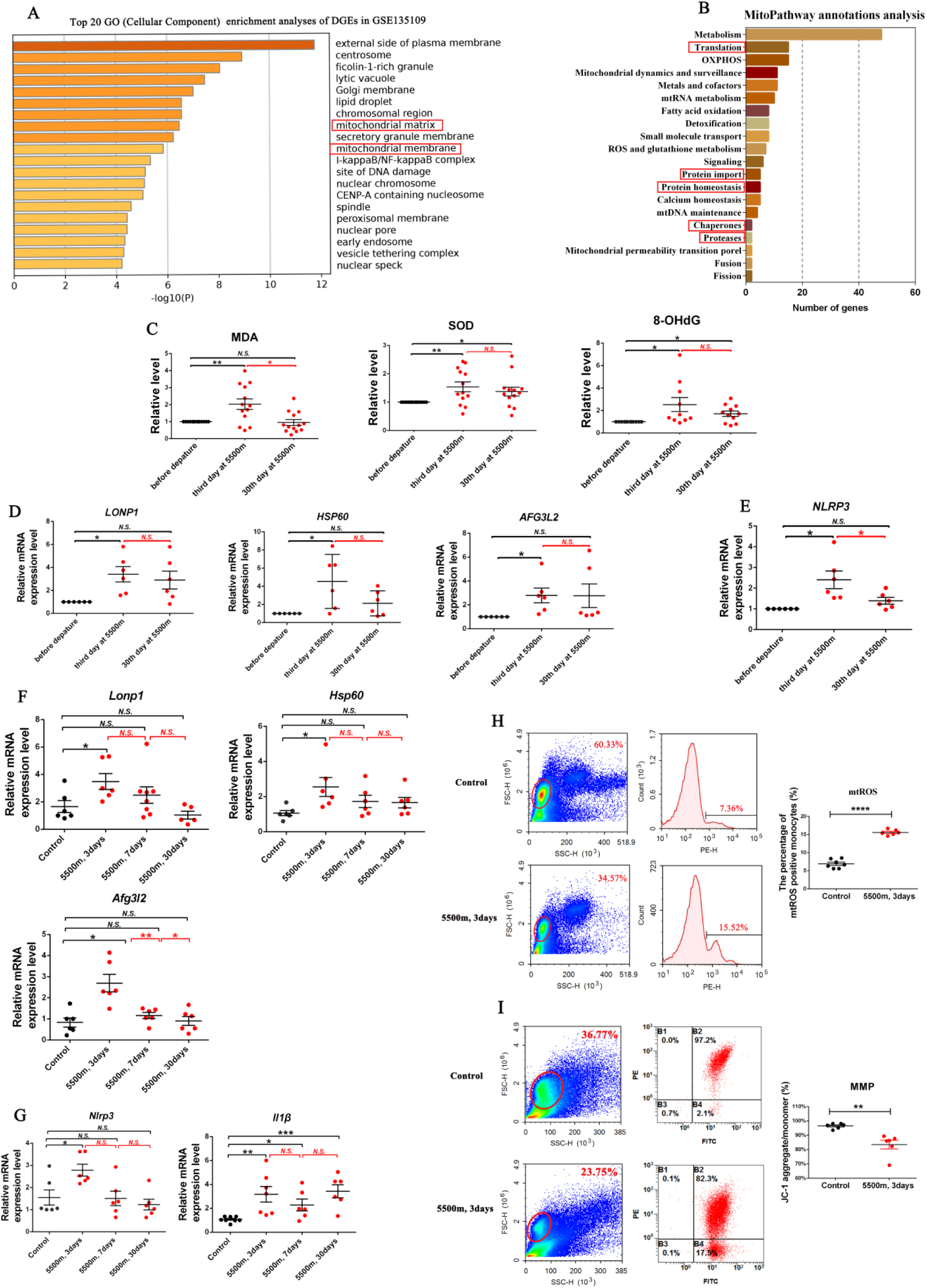
Hypobaric hypoxia induced the dynamic mitochondrial stress and mito-inflammation in PBMCs from human subjects and mouse model studies during 3-day and 30-day exposure. **A and B**, Top20 Cellular Component GO enrichment (**A**) and mitochondrial pathways annotations analyses (**B**) of DEGs in the human peripheral leukocytes between low-altitude group (stayed at sea level; n=3) and acute high-altitude group (stayed at the Qinghai-Tibet Plateau for 3 and 7 days; n=6) Red frames indicate the genes located in mitochondria (A) and the pathways related to mitochondrial proteins balance and homeostasis (B) (source data were obtained from GSE135109). **C**, Dynamic expression levels of oxidative stress related factors (SOD, MDA and 8-OHdG) in human plasma during the process of altitude climbing, by Luminex assay (n=20; normalized for nonspecific binding). **D and E**, Dynamic mRNA expression levels of genes involved mitochondrial unfolded protein response (UPR^mt^) (LONP1, HSP60, and AFG3L2) (**D**) and mitochondria-related inflammasome signaling (NLRP3) (**E**) in human PBMCs during the process of altitude climbing (n=8). **F and G**, Relative mRNA expression levels of genes involved UPR^mt^ (LONP1, HSP60, and AFG3L2) (**F**) and inflammasome signaling (NLRP3 and IL1β) (**G**) in the PBMCs of mice subjected into hypobaric chamber at the simulated altitude of 5500m for 3, 7 and 30 days respectively, by RT-qPCR analysis (n=6). **H and I**, Mitochondrial ROS (**H**) and mitochondrial membrane potential (**I**) levels in the circulating monocytes of mice subjected to the simulated altitude of 5500m for 3 days, by flow cytometric analysis (n=6). **C-I**, Data represented as mean± SEM. *P<0.05, **P<0.01, *** P<0.001, **** P<0.0001. Paired Student’s 2-tailed t test (C-D) and unpaired Student’s 2-tailed t test with Welch’s correction (F-I) were used to determine statistical significance.

In animal studies, mice were subjected into the hypobaric chamber at the simulated altitude of 5500 m for 3 days, 7 days and 30 days, respectively. The markers of mitochondrial stress response (*Lonp1*, *Afg3l2*, and *Hsp60*) and mito-inflammation (*Nlrp3* and *IL-1β*) in PBMCs were investigated by RT-qPCR (Fig. 2F and Fig. 2G). As compared with control group, the expression levels of genes involved mitochondrial stress and NLRP3 signaling in PBMCs significantly increased at 5500 m for 3 days and returned to the control levels on the 7-day or 30-day exposure of 5500 m. However, the level of *IL-1β* remained activation from 3-day to 30-day exposure of 5500 m (Fig. 2F and 2G). In addition, flow cytometry gating strategy was used to sort the monocytes in PBMCs (Fig. 2H and Fig. 2I, left) and then fluorescence probs were used to label the mitochondrial reactive oxygen species (mtROS) and mitochondrial membrane potential (MMP) in monocytes (Fig. 2H and 2I, right), which indicated that mtROS significantly increased but MMP decreased in the mouse circulating monocytes at 5500m for 3 days. Together, human and animal studies suggested that mitochondrial stress and mito-inflammation in PBMCs were obviously triggered in response to acute exposure of hyperbaric hypoxia, but gradually faded in the prolonged exposure of high altitude.

### 3. Hypobaric hypoxia promoted BMAL1 activation in the process of mitochondrial stress and mito-inflammation of monocytes

To identify the potential checkpoints related to mitochondria and inflammation in response to the acute hypobaric hypoxia, spearman correlation analyses based on GSE196728 was conducted and suggested that the increased abundance of monocytes in human whole blood cells were positively correlated with the activation of circadian gene *Bmal1* when exposed at 5100 m (Fig. 3A). Furthermore, GO annotations analysis based on GSE148510 confirmed that there was a significant proportion of DEGs located in the organelles and involved immune responses when *Bmal1* was deleted in bone marrow derived macrophages (BMDM) (Fig. S1E and Fig. 3B, red frames), which suggested that *Bmal1* could integrate the mitochondrial homeostasis and inflammatory response in PBMCs as reported previously^28^.

**Figure 3.**
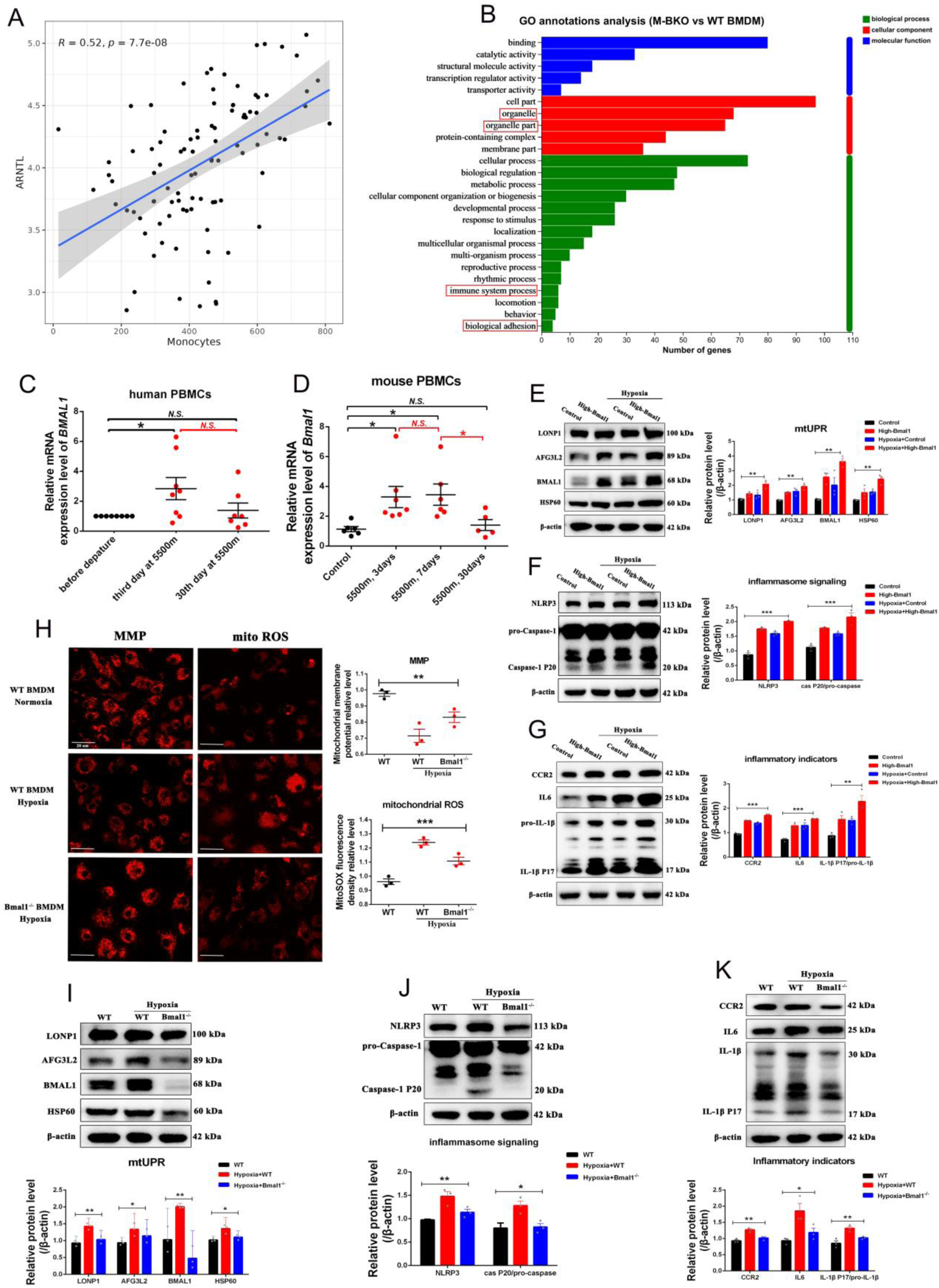
Hypobaric hypoxia promoted BMAL1 activation in the process of mitochondrial stress and mito-inflammation of monocytes. **A,** Spearman correlation analyses of *Bmal1* expression and monocytes abundance in human whole blood at La Rinconada (RI, Peru 5,100m) (source data were obtained from GSE196728). **B**, GO annotations analysis of DEGs in the Bone Marrow Derived Macrophages (BMDM) between WT mice and myeloid-specific *Bmal1* knock-out (M-BKO) mice. Red frames indicate the genes located in organelles and the pathways related to inflammation (n=3; source data were obtained from GSE148510). **C**, Dynamic mRNA expression of *Bmal1* in human PBMCs during the process of altitude climbing, by RT-qPCR analysis (n=8). **D**, Relative mRNA expression of *Bmal1* in PBMCs of mice subjected to simulated altitude of 5500m for 3, 7 and 30 days respectively, by RT-qPCR analysis (n=6). **E-G**, RAW264.7 cells were transfected with lentivirus to construct stable *Bmal1* overexpressed mouse monocytes, and then cultured under 21% oxygen normal or 1% oxygen hypoxic conditions for 24h. Relative proteins levels of genes involved UPR^mt^ (LONP1, HSP60, and AFG3L2) (**E**), mitochondria-related inflammasome signaling (NLRP3, pro-Caspase-1and Caspase-1 P20) (**F**) and inflammatory cytokines (IL6, IL-1βand CCR2) (**G**), by Western blotting analysis (n=3). **H-K**, *Bmal1* deficient mouse Bone Marrow Derived Macrophages (*Bmal1*^-/-^ BMDM) and wild-type BMDM (with normal expression of *Bmal1*) were isolated from the global *Bmal1* knock-out mice and WT mice respectively. **H**, Representative images and quantification of mitochondrial membrane potential (**left**) and mitochondrial ROS staining (**right**) in the *Bmal1*^-/-^ BMDM and WT BMDM under 1% oxygen hypoxic conditions for 24h (n=3); Scale bar = 20μm. **I-K**, Relative proteins levels of genes involved UPR^mt^ (LONP1, HSP60, and AFG3L2) (**I**), mitochondria-related inflammasome signaling (NLRP3, pro-Caspase-1and Caspase-1 P20) (**J**) and inflammatory cytokines (IL6, IL-1β and CCR2) (**K**) in the *Bmal1*^-/-^ BMDM and WT BMDM under 1% oxygen hypoxic conditions for 24h (n=3). **C-K**, Data represented as mean± SEM. *P<0.05, **P<0.01, *** P<0.001, **** P<0.0001. Unpaired Student’s 2-tailed t test with Welch’s correction (D), paired Student’s 2-tailed t test (C) and ordinary 1-way ANOVA followed by Tukey multiple comparisons (E-K) were used to determine statistical significance.

In the present work, the mRNA levels of *Bmal1* in human (Fig. 3C) and mouse PBMCs (Fig. 3D) significantly increased on 3-day exposure of 5500 m, whereas reduced to the control level on 30-day exposure of 5500 m, which was in consistent with the dynamic inflammatory response at different time points (Fig. 1G), indicating that acute exposure of hypobaric hypoxia induced the activation of *Bmal1*. In vitro study, *Bmal1* were overexpressed in the monocyte cell line RAW264.7 by lentivirus transfection (Fig. 3E-3G) and deleted in Bone Marrow Derived Macrophages (*Bmal1*^-/-^ BMDM) by isolating from the global *Bmal1* knock-out mice (Fig. 3H-3K), respectively. As shown by Western blotting, overexpression of *Bmal1* (High-Bmal1) significantly increased the marker expressions of mitochondrial stress (LONP1, AFG3L2 and HSP60, Fig. 3E), mito-inflammasome signaling (NLRP3, pro-Caspase-1 and Caspase-1 P20, Fig. 3F), and inflammatory response (IL6, MCP-1 and IL-1β, Fig. 3G) in RAW264.7 cells under 21% oxygen normal condition for 24 h. When exposed to 1% oxygen hypoxia, overexpression of *Bmal1* (High-Bmal1) furtherly increased the marker expressions of mitochondrial stress, mito-inflammasome signaling, and inflammatory response in RAW 264.7 cells as compared with that of control (Figure 3E-3G). In contrast, deletion of *Bmal1* (*Bmal1*^-/-^) not only obviously alleviated oxygen stress by reducing mitochondrial reactive oxygen species (mtROS) and increasing the mitochondrial membrane potential (MMP) (Fig. 3H), but also markedly decreased the marker expressions of mitochondrial stress (LONP1, AFG3L2 and HSP60, Fig. 3I), mito-inflammasome signaling (NLRP3, pro-Caspase-1 and Caspase-1 P20, Fig. 3J), and inflammatory response (IL6, MCP-1 and IL-1β, Fig. 3K) in BMDM when exposed to 1% oxygen hypoxia. Our In vivo and in vitro study indicated that activation of *Bmal1* aggravated mitochondrial stress and inflammatory response in monocytes under acute hypobaric hypoxia.

### 4. BMAL1 stabilized mitochondrial protein homeostasis by targeting Fis1 transcription in monocytes

Under environmental stress, mitochondrial fission and fusion are known to play critical roles in maintaining the integrity and homeostasis of mitochondria, and mitochondrial dynamics is a kind of important mitochondrial quality control mechanism for maintaining mitochondrial protein homeostasis. The Venn diagram of DEGs in GSE199947 and Human MitoCarta3.0 suggested that the pathways of mitochondrial dynamics and mitochondrial protein homeostasis changed most significantly in response to hypobaric hypoxia (Fig. 4A and Fig. S1F, red frames). In the present work, mRNA expression levels of both *Bmal1* and mitochondrial fission genes in PBMCs (*Fis1* and *Drp1*) significantly increased, whereas the mitochondrial fusion genes (*Mfn1*, *Mfn2* and *Opa1*) obviously decreased as compared with that of control mice when exposed to the simulated altitude of 5500 m for 3 days (Fig. 4B). To investigate the mitochondrial downstream target of *Bmal1*, the DNA fragments in mouse monocyte cell line RAW 264.7 were absorbed by magnetic beads coated with BMAL1 ChIP antibody and then analyzed by RT-PCR, which suggested that BMAL1 directly bond to the promoter region of *Fis1* and enhanced its transcription (Fig. 4C). In addition, deletion of *Bmal1* significantly decreased the mRNA (Fig. 4D) and protein expression levels of *Fis1* (Fig. 4E) in BMDM, whereas overexpression of *Bmal1* in RAW 264.7 cells significantly increased FIS1 protein expression under both normoxic and hypoxic conditions as compared with their relative control group (Fig. 4F). To investigate the role of *Bmal1* and its targeting Fis1 in mitochondrial morphology and function, the BMDM isolated from wild-type (WT) and *Bmal1*^-/-^ mice were cultured under 1% oxygen hypoxia for 24 h. As shown in Fig. 4G, hypoxia induced an abundance of punctate mitochondria with loss of filamentous in WT BMDM; however, deletion of *Bmal1* significantly recovered the population of rod-like mitochondria in the BMDM exposed to hypoxia. Consistently, hypobaric hypoxia significantly increased the mitochondrial fission proteins (DRP1 and FIS1) and decreased the mitochondrial fusion proteins (OPA1, MFN1 and MFN2) in the WT BMDM; while deletion of *Bmal1* restored the protein expressions of mitochondrial fission and fusion proteins to the control level (Fig. 4H). Together, these results suggested activation of BMAL1 targeted *Fis1* transcription and then regulating mitochondrial integrity and homeostasis under hypoxia stress.

**Figure 4.**
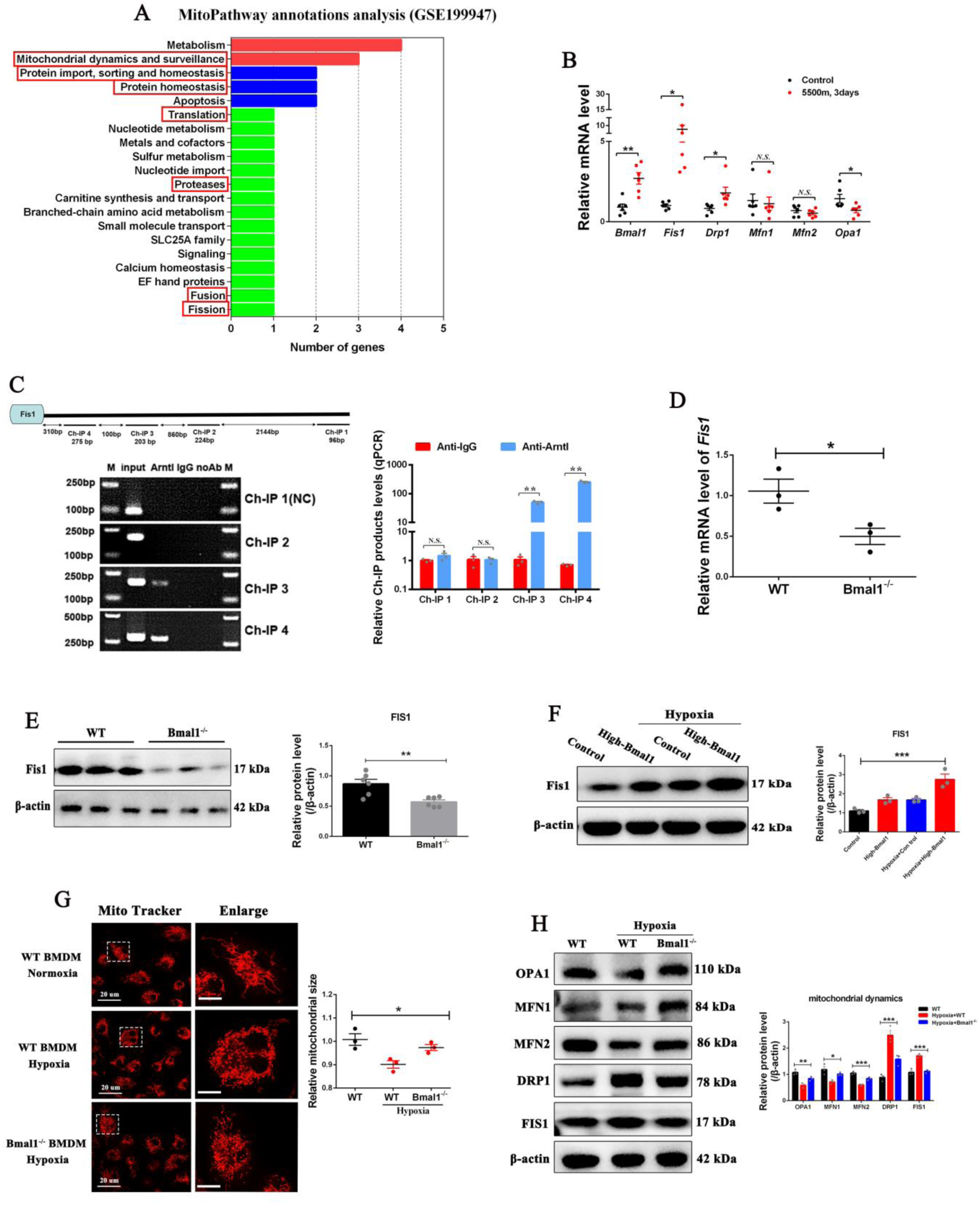
BMAL1 stabilized mitochondrial protein homeostasis by targeting Fis1 transcription in monocytes. **A**, Mitochondrial pathways annotations analysis of DEGs in the human monocytic THP-1 cells between normoxia group (n=3) and hypoxia group (under 1% oxygen hypoxic condition for 8 and 72h; n=6). Red frames indicate the pathways of mitochondrial dynamics and protein homeostasis. (source data were obtained from GSE199947). **B**, Relative mRNA expression levels of *Bmal1*, mitochondrial fission genes (*Fis1* and *Drp1*) and mitochondrial fusion genes (*Mfn1*, *Mfn2* and *Opa1*) in the PBMCs of mice exposed to simulated altitude of 5500m for 3 days, by RT-qPCR analysis (n=6). **C**, Chromatin immunoprecipitation (ChIP) assays for BMAL1 enrichment to the *Fis1* promoter on chromatin prepared from RAW264.7 cells. RT-PCR (**left**) and RT-qPCR (**right**) analyses of ChIP products with 4 pairs of amplification primers (including a pair of control primers) (n=3; IgG immunoprecipitation was used as control). **D, G and H,** *Bmal1* deficient (*Bmal1*^-/-^) BMDM and wild-type (WT; with normal expression of *Bmal1*) BMDM were isolated from *Bmal1*^-/-^ mice and WT mice respectively. **D**, Relative mRNA expression level of *Fis1* in WT BMDM and *Bmal1*^-/-^ BMDM, by RT-qPCR analysis (n=3). **E**, Relative protein level of FIS1 in WT BMDM and *Bmal1*^-/-^ BMDM, by Western blotting analysis (n=3). **F**, Relative protein level of FIS1 in *Bmal1* overexpressed RAW264.7 cells cultured under 21% oxygen normal or 1% oxygen hypoxic conditions for 24h respectively, by Western blotting analysis (n=3). **G**, Mitochondrial sizes of *Bmal1*^-/-^ BMDM and WT BMDM under 1% oxygen hypoxic conditions for 24h, by Mito-Tracker staining (n=3); Scale bar = 20μm. **H**, Relative levels of mitochondrial fission proteins (FIS1 and DRP1) and mitochondrial fusion proteins (MFN1, MFN2 and OPA1) in *Bmal1*^-/-^ BMDM and WT BMDM under 1% oxygen hypoxic conditions for 24h, by Western blotting analysis (n=3). **B-H**, Data represented as mean± SEM. Significance was determined by ordinary 1-way ANOVA with Tukey multiple comparisons. Unpaired Student’s 2-tailed t test with Welch’s correction (B-E) and ordinary 1-way ANOVA followed by Tukey multiple comparisons (I, F-H) were used to determine statistical significance.

### 5. Activation of *Bmal1* drove mitochondrial stress and mito-inflammation in monocytes by promoting Fis1-mediated mitochondrial fission in response to hypoxia

The balance between mitochondrial fission and fusion is essential for mitochondrial integrity and excessive mitochondrial fission leads to mitochondrial fragmentation and thus activates stress signaling. Mitochondrial division inhibitor-1 (Mdivi-1) was used to investigate the role of mitochondrial fission in the hypoxia-induced mitochondrial stress and mito-inflammation in monocytes. As shown in Fig. 5A, Mdivi-1 significantly inhibited mitochondrial fission in in dose and time dependent manner. Under normoxic condition, overexpression of *Bmal1* induced an obvious mitochondrial dysfunction characterized with the reduced MMP and increased mitochondrial ROS in RAW264.7 cells. In addition, *Bmal1* overexpression significantly induced mitochondrial stress with increased expression of LONP1, AFG3L2, and HSP60 (Fig. 5C), and triggered mito-inflammatory signaling with the enhanced expression of NLRP3, pro-Caspase-1, and Caspase-1 P20 (Fig. 5D), and then induced the inflammatory response with the increased expression of IL6, CCR2, and IL-1β (Fig. 5E) in RAW264.7 cells. However, inhibition of mitochondrial fission by 50 uM Mdivi-1 remarkedly reversed the *Bmal1* overexpression-induced MMP depolarization and mtROS production (Fig. 5B), and alleviated the *Bmal1* overexpression-induced mitochondrial stress, mito-inflammation, and inflammatory response (Fig. 5C-5E) in RAW264.7 cells. When exposed to hypoxia, the mitochondrial dysfunctions (Fig. 5B), mitochondrial stress (Fig. 5C), mito-inflammatory signaling (Fig. 5D), and inflammatory response (Fig. 5E), which were induced by hypoxia and furtherly aggregated by *Bmal1* overexpression, were significantly alleviated by the treatment of Mdivi-1, respectively. Together, these results suggested that *Bmal1* triggered mitochondrial stress and mito-inflammation by promoting mitochondrial fission in monocytes when exposed to hypoxia.

**Figure 5.**
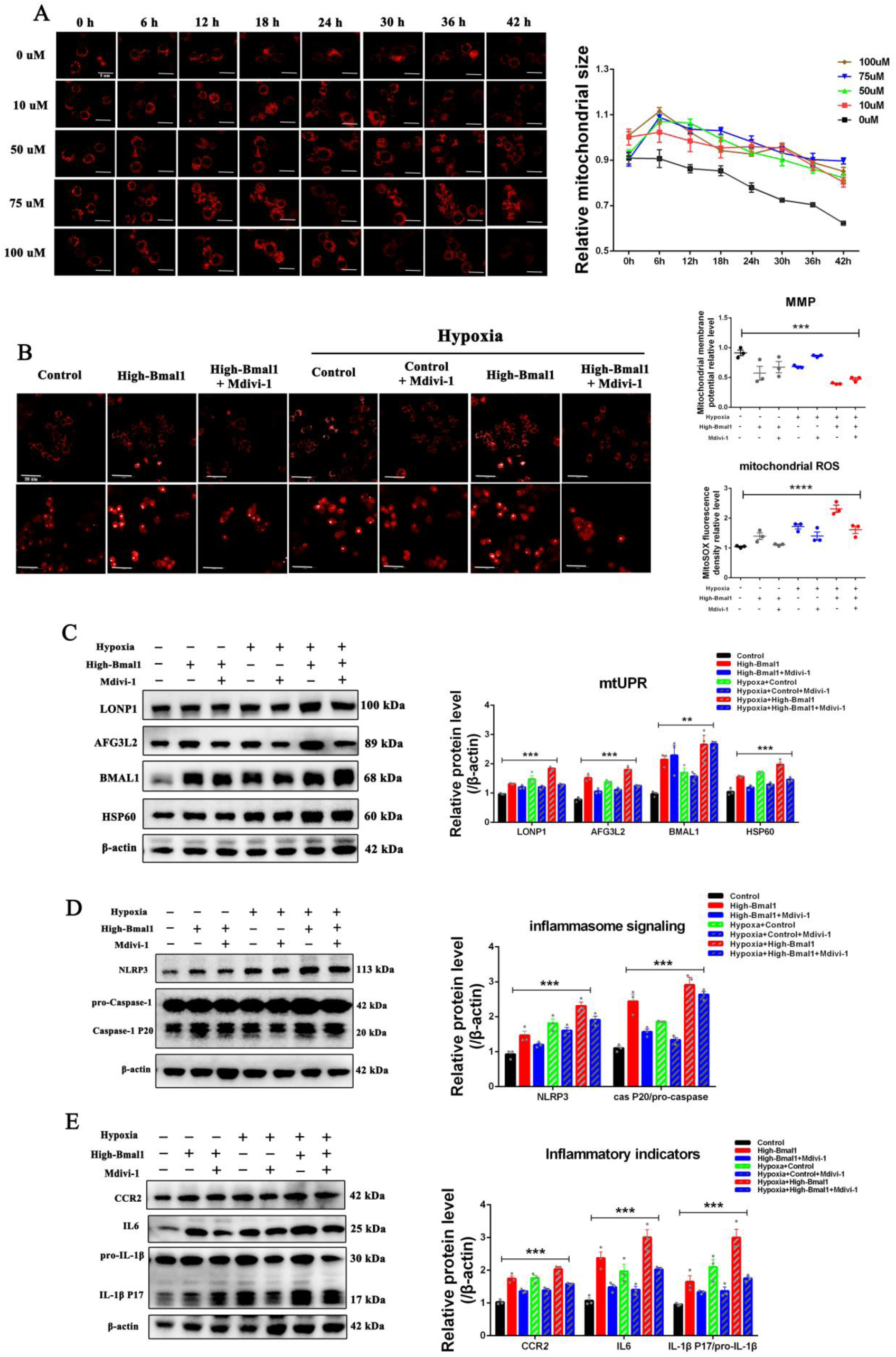
Activation of *Bmal1* drove mitochondrial stress and mito-inflammation by promoting Fis1-me dieted mitochondrial fission in monocytes under hypoxia. **A**, RAW264.7 cells were treated with mitochondrial fission inhibitor (Mdivi-1) in the concentrations of 0uM, 10uM, 50uM, 75uM and 100uM, respectively. The dynamic changes of mitochondrial sizes in the RAW264.7 cells following 0 to 42h Mdivi-1 incubation, by Mito-Tracker staining (n=3); Scale bar = 5μm. **B-E**, *Bmal1* overexpressed RAW264.7 cells were treated with mitochondrial fission inhibitor (50uM Mdivi-1) and then cultured 21% oxygen normal or 1% oxygen hypoxic conditions for 24h. **B**, Representative images and quantification of mitochondrial membrane potential (**upper**) and mitochondrial ROS staining (**down**) (n=3); Scale bar = 50μm. **C-E**, Relative proteins levels of genes involved UPR^mt^ (LONP1, HSP60, and AFG3L2) (**C**), mitochondria-related inflammasome signaling (NLRP3, pro-Caspase-1 and Caspase P20) (**D**) and inflammatory cytokines (IL6, IL-1βand CCR2) (**E**), by Western blotting analysis (n=3); **A-E**, Data represented as mean± SEM. Significance was determined by ordinary 1-way ANOVA with Tukey multiple comparisons.

### 6. Transcription factor BHLHE40 triggered *Bmal1* activation in monocytes under hypoxia

So far, we have clarified that the transcriptional activation of *Bmal1* in monocytes is a driving factor for initiating the mitochondrial stress and mito-inflammation under hypoxia, but limited study ever reported which factors could trigger *Bmal1* transcription. Histone modifications and transcriptional factors (TFs) have been previously reported to be the two important mechanisms accounting for the gene transcriptional activation, hence we integrated transcriptomics and bioinformatic analyses (Fig. 6A), aiming to find out which types of histone modifications and TFs could account for the transcriptional activation of *Bmal1* in monocytes under hypoxia. Based on ChIPBase v3.0 (https://rnasysu.com/chipbase3/index.php), 29 types of histone modifications were predicted to regulate *Bmal1* transcription (Table 1), among which the methylated modification in lysine 4 of histone H3 (H3K4me1/2/3) may be mostly responsible for the hypoxia signaling, since KDM5 histone demethylases family are a group of oxygen-dependent dioxygenase, which could target H3K4me1/2/3 to regulate genes expression. In the present work, on the one hand, we confirmed that hypoxia significantly increased the protein levels of H34Kme3 in PBMCs of mice subjected to the simulated altitude at 5500 m for 3 days (Fig. 6B) and in the RAW264.7 cells cultured under 1% oxygen hypoxic conditions for 12 h and 24 h (Fig. 6C), respectively. However, inhibition of KDM5 demethylases by CPI-455 (the KDM5 demethylases inhibitor, Fig. 6D) or siRNAs transfection against *Kdm5a* and *Kdm5c* (Fig. 6E) did not significantly affect the mRNA expression of *Bmal1*, suggesting that the KDM5 demethylases-mediated H3K4me1/2/3 did not trigger the *Bmal1* transcription when exposed to hypoxia. On the other hand, based on the JASPAR database, we predicted 68 upstream transcription factors (TFs) which potentially bind to the promoter regions and regulatory sites of *Bmal1* (Table 2). In addition, through taking intersection of the database of GSE135109 (mRNA expression profiles of peripheral leukocytes in human subjects under hypobaric hypoxia) and GSE199947 (mRNA expression profiles of THP-1 monocytes under hypoxia), we identified 16 common DEGs in monocytes when exposed to hypoxia, and the 16 DEGs were enriched in the pathway of regulation of DNA-binding transcription factor activity (Fig. 6F and 6G). By Ingenuity Pathway Analysis (IPA), the basic helix-loop-helix family member e40 (BHLHE40) was screened out from the 234 candidate factors (Table 3) which act on the 68 upstream TFs of *Bmal1* (Table 2) (Fig. S3A and Fig. 6H). BHLHE40 is an immediate-early response gene in macrophages, which has been reported to a kind of HIF-1α targeting transcriptional factor (Fig. S3B). The present work confirmed that inhibition of *Bhlhe40* by siRNAs transfection significantly decreased the mRNA expression of *Bmal1* in RAW264.7 cells (Fig. 6I).

**Figure 6.**
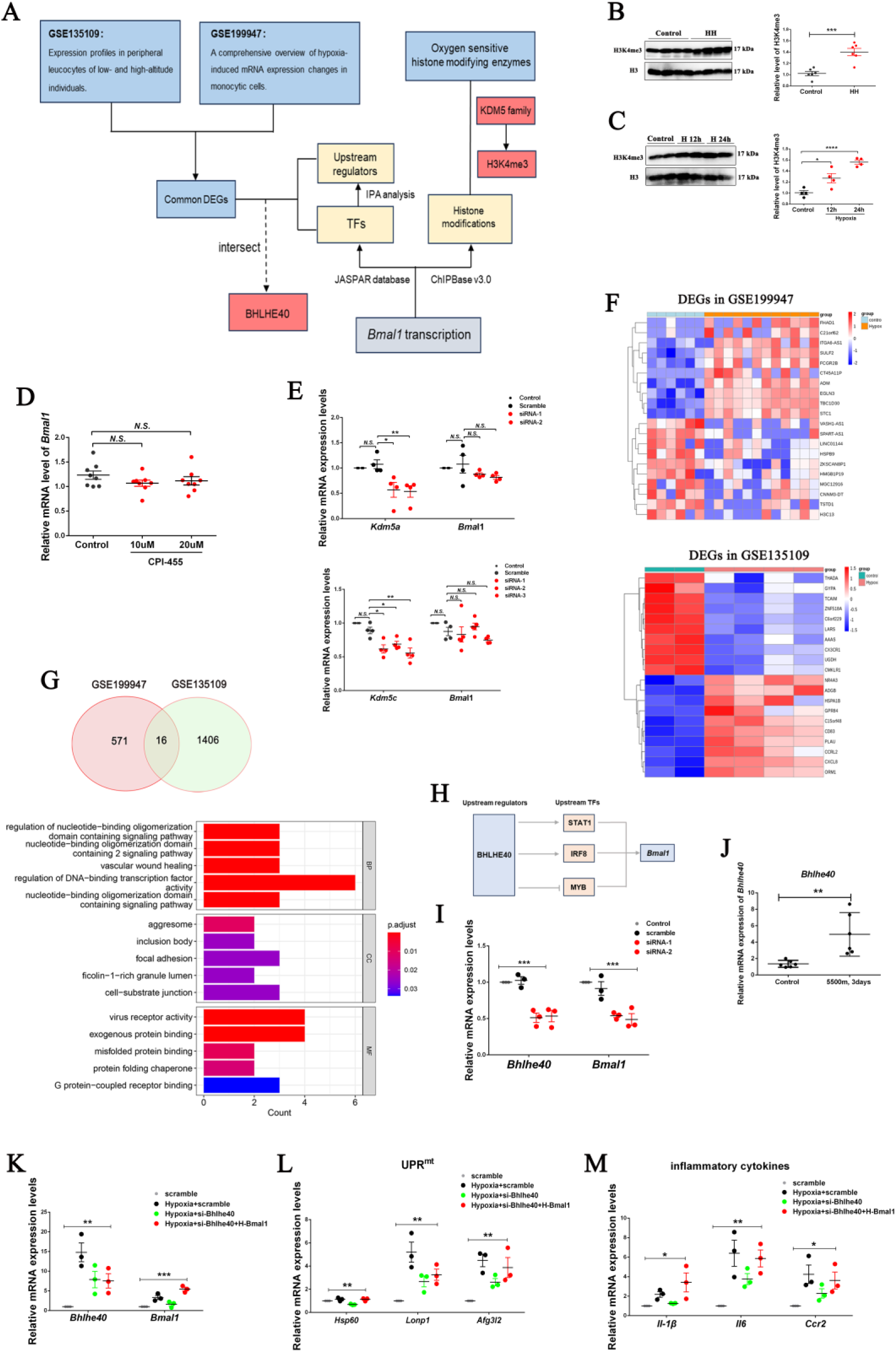
Transcription factor BHLHE40 triggered *Bmal1* activation in monocytes under hypoxia. **A,** Flow chart of bioinformatics analysis to find out the potential regulators for the *Bmal1* transcriptional activation under hypoxia. **B**, Relative protein level of H34Kme3 in the PBMC of mice exposed to simulated altitude of 5500m for 3 days by hypobaric chamber, by Western blotting analysis (n=6). **C**, Relative protein level of H34Kme3 in RAW264.7 cells cultured under 1% oxygen hypoxic conditions for 12h and 24h respectively, by Western blotting analysis (n=3). **D**, Relative mRNA expression level of *Bmal1* in RAW264.7 cells treated with KDM5 family inhibitor (CPI-455) in the concentrations of 10 uM and 20 uM for 24h respectively, by RT-qPCR analysis (n=3). **E**, Relative mRNA expression level of *Bmal1* in RAW264.7 cells transfected with *Kdm5a* (upper) and *Kdm5c* (down) siRNAs, by RT-qPCR analysis (n=3). **F**, Heatmap of DEGs in the human monocytic THP-1 cells between normoxia group (n=3) and hypoxia group (under 1% oxygen hypoxic condition for 8 and 72h (n=6) (source data were obtained from GSE199947; **upper**) and heatmap of DEGs in the human peripheral leukocytes between low-altitude group (stayed at sea level; n=3) and high-altitude group (stayed at the Qinghai-Tibet Plateau for 3 and 7 days; n=6) (source data were obtained from GSE135109; **down**). Various color intensities indicate the expression levels: bottom expressions are shown by blue and top by red. **G,** Venn diagram of DEGs intersection between GSE135109 and GSE199947 **(upper),** and GO pathway enrichment analysis of the 16 intersected genes **(down). H,** Diagram of the predicted regulatory pathways of BHLHE40 on *Bmal1* transcription. **I,** Relative mRNA expression level of *Bmal1* in RAW264.7 cells transfected with *Bhlhe40* siRNAs, by RT-qPCR analysis (n=3)**. J,** Relative mRNA expression level of *Bhlhe40* in the PBMCs of mice subjected to the simulated altitude of 5500m for 3 days, by RT-qPCR analysis (n=6)**. K-M,** *Bmal1* overexpressed RAW264.7 cells were transfected with siRNAs to inhibit the *Bhlhe40* expression, and then 1% oxygen hypoxic condition. Relative mRNA expression levels of *Bmal1* **(K),** UPR^mt^ genes (*Lonp1*, *Hsp60*, and *Afg3l2*) **(L),** and inflammatory cytokines (*Il6*, *Mcp1*, and *Il1β*) **(M)**, by RT-qPCR analysis (n=3)**. *P<0.05, **P<0.01, *** P<0.001, **** P<0.0001.** Unpaired Student’s 2-tailed t test with Welch’s correction (B-E, J) and ordinary 1-way ANOVA followed by Tukey multiple comparisons (I, K-M) were used to determine statistical significance.

**Table 1.** The prediction of histone modifications in regulatory regions of *Bmal1*.

| Histone modification | Number of samples found |  | Number of binding sites |  |
| --- | --- | --- | --- | --- |
|  | upstream | downstream | upstream | downstream |
| H1.0 | 0 | 1 | 0 | 1 |
| H1.4 | 1 | 2 | 1 | 2 |
| H2A | 5 | 3 | 5 | 3 |
| H2AFZ | 14 | 13 | 14 | 14 |
| H2AK9ac | 1 | 1 | 1 | 2 |
| H2AZ | 5 | 1 | 6 | 1 |
| H2BK120ac | 1 | 1 | 1 | 1 |
| H2BK5ac | 2 | 2 | 2 | 2 |
| H3F3A | 4 | 1 | 4 | 2 |
| H3K14ac | 0 | 3 | 0 | 4 |
| H3K18ac | 3 | 10 | 3 | 13 |
| H3K23ac | 2 | 2 | 2 | 4 |
| H3K23me2 | 1 | 2 | 1 | 3 |
| H3K27ac | 10 | 11 | 10 | 11 |
| H3K27me1 | 0 | 1 | 0 | 1 |
| H3K4me1 | 3 | 1 | 3 | 1 |
| H3K4me2 | 20 | 22 | 20 | 22 |
| H3K4me3 | 9 | 37 | 9 | 38 |
| H3K4me3B | 1 | 1 | 1 | 1 |
| H3K9ac | 11 | 12 | 11 | 13 |
| H3K9me3 | 1 | 0 | 1 | 0 |
| H3ac | 2 | 2 | 2 | 4 |
| H4 | 0 | 1 | 0 | 1 |
| H4K20me1 | 0 | 3 | 0 | 3 |
| H4K20me3 | 0 | 1 | 0 | 1 |
| H4K5ac | 1 | 2 | 1 | 2 |
| H4K8ac | 2 | 3 | 3 | 5 |
| H4K91ac | 1 | 3 | 1 | 3 |
| H4ac | 1 | 3 | 1 | 3 |

<sup>a</sup> Based on ChIPBase v3.0 (https://rnasysu.com/chipbase3/index.php), 29 types of histone modifications were predicted to occur in the regulatory regions of *Bmall*.

**Table 2.** The prediction of upstream transcriptional factors of *Bmal1*.

| **Symbol** | **Symbol** | **Symbol** | **Symbol** |
| --- | --- | --- | --- |
| GATA3 | FOXD2 | IRF3 | KLF16 |
| TRPS1 | BCL11B | IRF8 | KLF11 |
| NR2F2 | HSF1 | IRF9 | KLF15 |
| MYOD1 | HSF2 | CDX1 | SP8 |
| SNAI2 | HSF4 | CDX2 | SP9 |
| SNAI3 | REST | HOXA10 | PATZ1 |
| MYB | ZIC1 | CDX4 | SP1 |
| RFX4 | NR2F2 | HOXD9 | KLF1 |
| ONECUT1 | NR2F6 | ZNF701 | SP2 |
| ZNF24 | IRF2 | TEAD1 | SP4 |
| NFIC | IRF4 | MEIS1 | KLF14 |
| ZNF549 | IRF1 | LHX2 | KLF12 |
| MGX | ZKSCAN1 | KLF9 | KLF10 |
| EVX1 | NR1H2 | ZNF281 | KLF7 |
| FOXD3 | NR4A1 | SP3 | WT1 |
| FOXJ3 | STAT2 | KLF5 | ZNF740 |
| FOXC2 | STAT1 | KLF16 | RREB1 |

<sup>a</sup> Based on the JASPAR database, 68 upstream transcription factors were predicted to act on the upstream 2000bp promoter region of *Bmal1*.

**Table 3.**
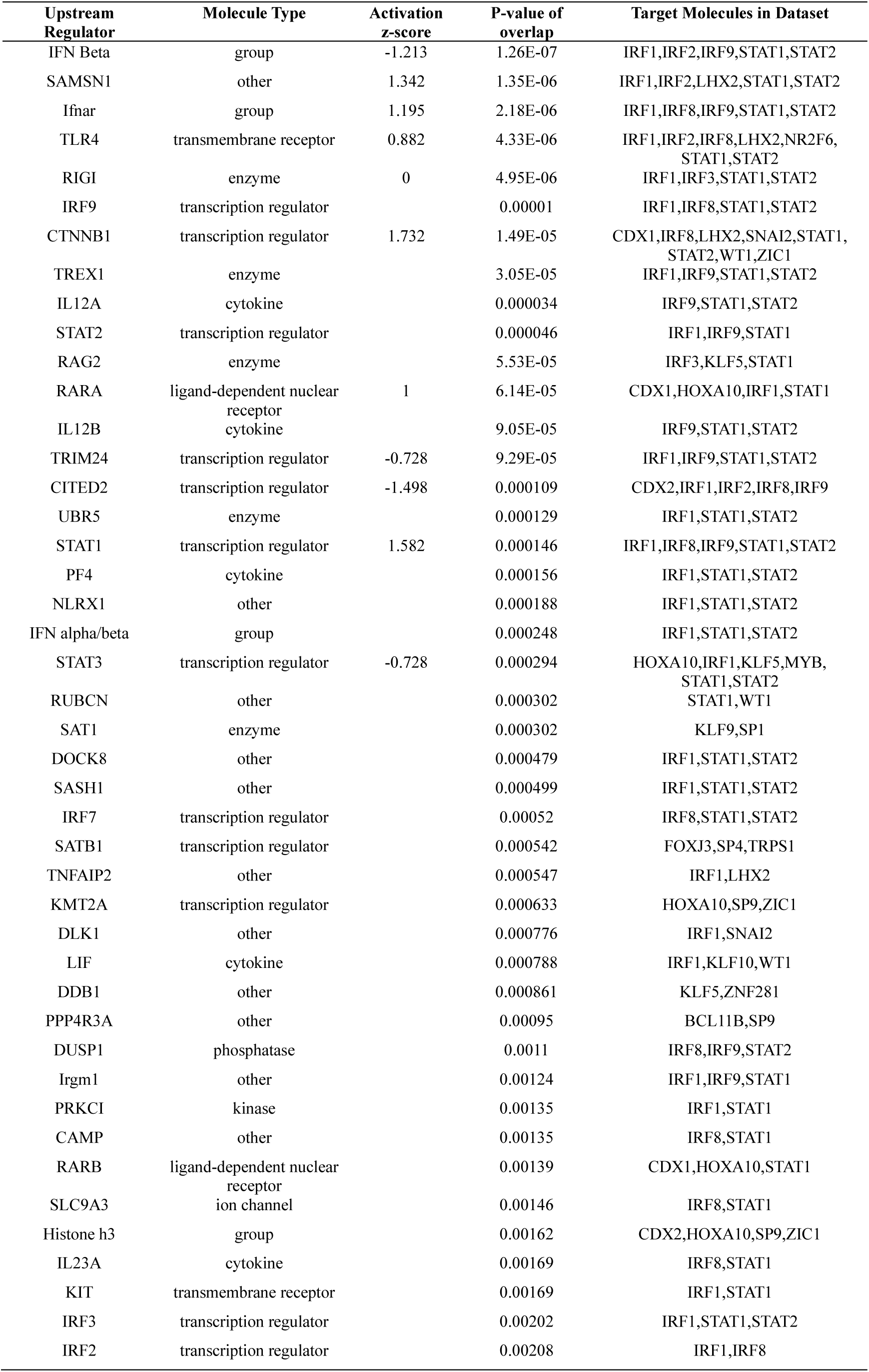
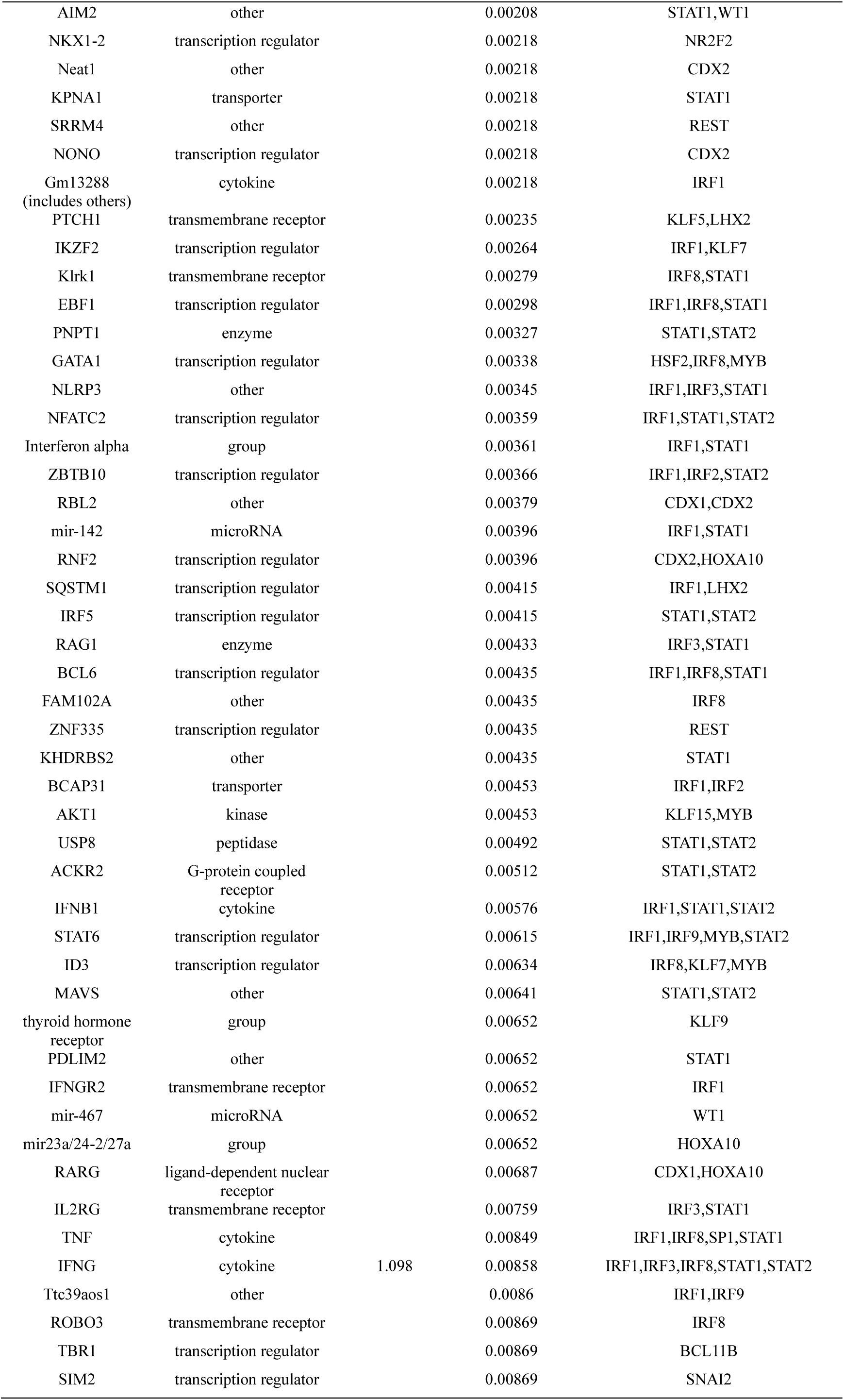
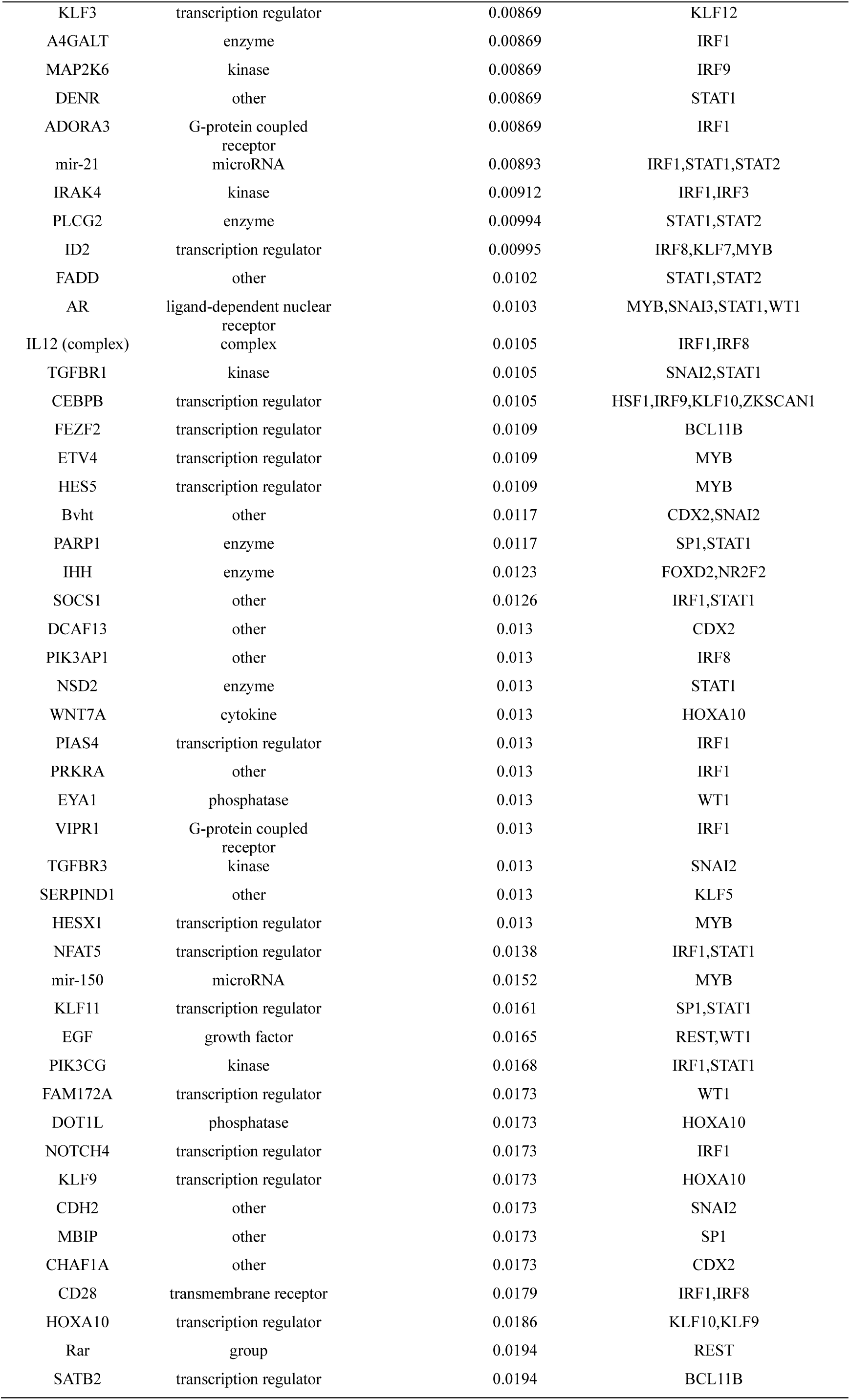
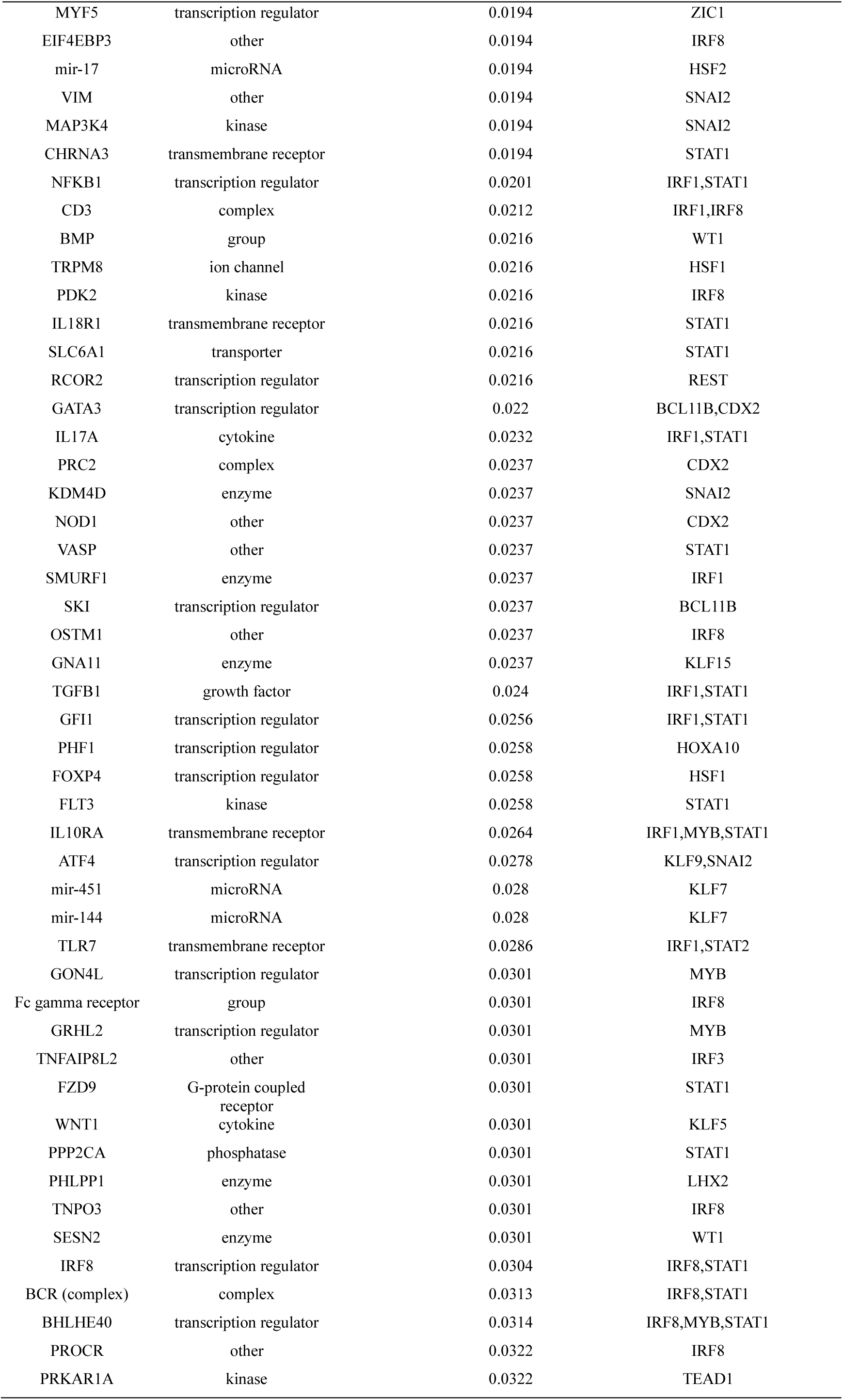
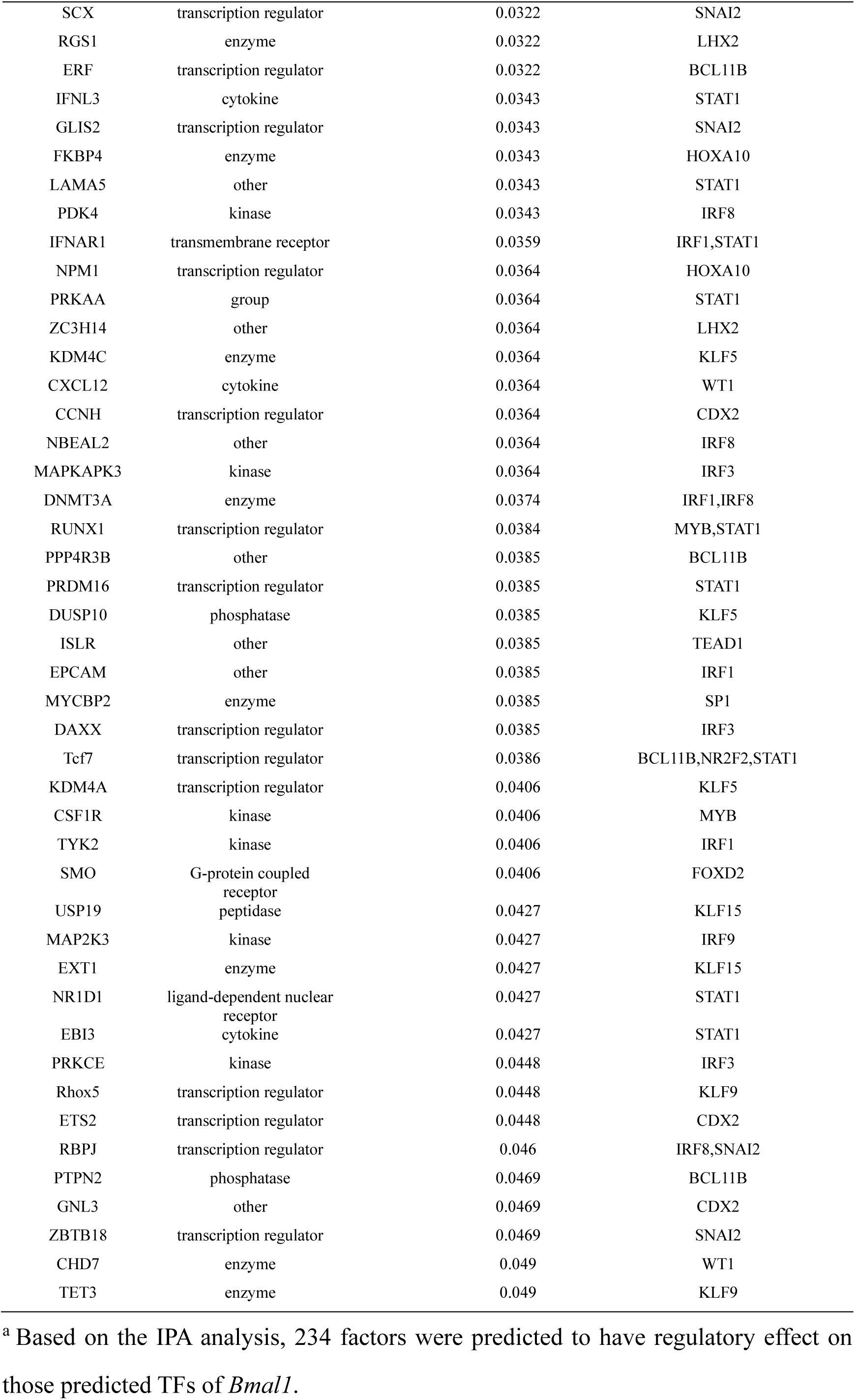
The prediction of upstream regulators of TFs targeting *Bmal1*.

When exposed to hypoxia, mRNA expression of *Bhlhe40* was found to be significantly increased in PBMCs of mice (Fig. 6J) and RAW264.7 cells (Fig. 6K), respectively. Inhibition of *Bhlhe40* by siRNAs transfection significantly alleviated the hypoxia-induced the transcription of *Bmal1* (Fig. 6K), the activation of mitochondrial stress (*Lonp1*, *Afg3l2*, and *Hsp60*, Fig. 6L), and the inflammatory response (*Il6*, *Ccr2*, and *Il1β*, Fig. 6M) in RAW264.7 cells. In addition, overexpression of *Bmal1* did not affect the mRNA expression of *Bhlhe40* (Fig. 6K), but significantly reversed the blocking effect of *Bhlhe40* in the activation of mitochondrial stress (*Lonp1*, *Afg3l2*, and *Hsp60*, Fig. 6L), and the inflammatory response (*Il6*, *Ccr2*, and *Il1β*, Fig. 6M) in RAW264.7 cells. These results suggested BHLHE40 was an upstream signaling of *Bmal1* transcriptional activation, and thus to elicit mitochondrial stress and inflammatory response in the monocytes under hypoxia.

### 7. Myeloid-specific *Bmal1* knock-out alleviated the systemic circulating inflammation and the local inflammatory microenvironment of pulmonary vasculature under hypobaric hypoxia

Inflammatory monocyte is a kind of significant executors of systemic inflammation and vascular inflammation. In the present work, we successfully generated the myeloid-specific *Bmal1* knock-out mice (M-BKO, *Bmal1*^f/f^ crossed to Lyz2-Cre) and exposed the M-BKO mice to the simulated altitude of 5500 m for 3 days. Knock-out of myeloid-specific *Bmal1* not only significantly reduced the ratio of inflammatory (Ly6C^Hi^) monocytes (Fig. 7A), but also obviously decreased the mRNA expressions of inflammatory cytokines in PBMCs (*Il6*, *Ccr2*, and *Il1β*, Fig. 7B) and the protein expressions of inflammatory cytokines in plasma (IL6, MCP-1, and Il1β, Fig. 7C) as compared with that of *Bmal1*^f/f^ mice (the wild-type control).

**Figure 7.**
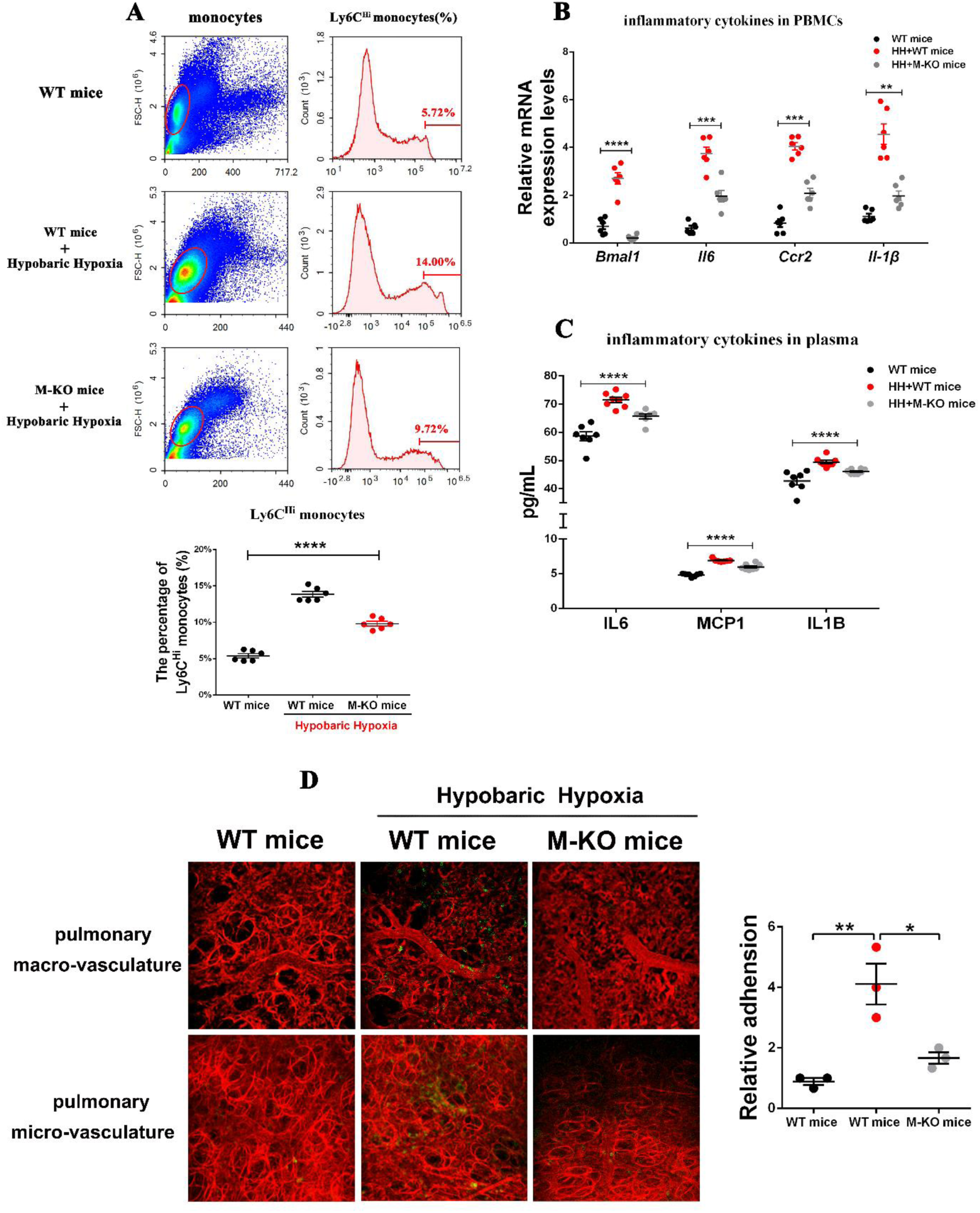
Myeloid-specific *Bmal1* knock-out alleviated the systemic inflammation and the local inflammatory microenvironment of pulmonary vasculature under hypobaric hypoxia. A-D, Myeloid-specific *Bmal1* knock-out mice were exposed to simulated altitude of 5500m for 3 days by hypobaric chamber. **A**, Inflammatory (Ly6C^Hi^) monocytes ratio in mouse circulating monocytes, by flow cytometric analysis (n=6-8). **B**, Relative mRNA expression levels of inflammatory cytokines (IL6, IL-1β, and CCR2) in mouse PBMCs, by RT-qPCR analysis (n=6-8). **C**, Protein levels of inflammatory cytokines (IL6, IL-1β, and MCP-1) in mouse plasma, by ELISA assay (n=6-8). **D**, In vivo imaging of inflammatory cells (CD11b^+^ cells) infiltrating into pulmonary vasculature, by Multichannel fluorescence intravital microscopy (MFIM) (red, angiography Evans blue; green, CD11b^+^ cells). **A-D**, Data represented as mean± SEM. *P<0.05, **P<0.01, *** P<0.001, **** P<0.0001. Statistical significance was determined by Ordinary 1-way ANOVA followed with Tukey multiple comparisons.

To observe the inflammatory cells infiltration into vasculature in real-time, WT and M-BKO mice were injected with low-dose CD11b^+^ fluorescent antibody to label the inflammatory cells in vivo, and then the interactions between inflammatory cells and pulmonary vasculature were visualized with multichannel fluorescence intravital microscopy (MFIM). As shown in Fig. 7D and Video S1-S3, exposure to acute hypobaric hypoxia induced obvious adhesion and infiltration of inflammatory cells into pulmonary macro- and micro-vasculatures. However, the levels of inflammatory cells adhesion and infiltration into vasculature were significantly alleviated in the M-BKO mice under acute hypobaric hypoxia. Together, these results suggested that myeloid-specific *Bmal1* knock-out reduced inflammatory signaling in the monocytes, and furtherly alleviated the inflammation at the circulating and vascular tissues levels under acute hypobaric hypoxia.

## Discussion

The major and novel findings of this study are as follows: 1) Hypobaric hypoxia induced a dynamic inflammatory response of PBMCs from human subjects and mouse models during 3-day and 30-day exposure. 2) Hypoxic stress triggered the mitochondrial unfolded protein response (mitochondrial stress) and then induced the mito-inflammation (NLRP3 inflammasome) in monocytes. 3) Activation of *Bmal1* drove mitochondrial stress and mito-inflammation by promoting Fis1-mediated mitochondrial fission in monocytes under hypoxia. 4) BHLHE40, a stress-responsive transcription factor directly targeted by HIF-1α, stimulated *Bmal1* transcription in monocytes under hypobaric hypoxia. 5) Myeloid-specific *Bmal1* deletion alleviated systemic circulating inflammation and vascular inflammation under acute hypobaric hypoxia.

### Monocytes in PBMCs were more sensitive and contributed promptly to circulating inflammation in response to acute hypobaric hypoxia

As altitude increases, the decreased barometric pressure causes a reduction of oxygen partial pressure, resulting in a hypobaric hypoxic stress and leading to a range of altitude-related illnesses for individuals climbing, such as AMS, HAPE and HACE^33, 34^, as well as the exacerbation of pre-existing cardiovascular, pulmonary, gastrointestinal diseases and infections^3^. Previous studies have highlighted the critical role of hypoxia-induced inflammation in the development of these altitude-related issues^6–8, 9, 10^. It has been observed that increased leukocytes and inflammatory mediators in bronchoalveolar fluid typically manifest in the later stages of HAPE^6, 35^. In the present work, we found a significant increase of inflammatory cytokines (IL-1β, MCP-1, and IL-6) in circulating plasma on the third day and a continuous rise on the 30^th^ day at the altitude of 5500 m (Fig. 1A and 1B). It is noted that the severity of high-altitude illnesses is obviously influenced by exposure time for certain physiological responses may gradually diminish with the prolonged exposure durations^33, 36^, including the enhanced sympathetic activity and hypoxic pulmonary vasoconstriction. However, our present work indicated that inflammatory cytokines in the circulating plasma consistently increased in response to high-altitude hypoxia.

As central players of the innate immune system, monocytes exert an important function by orchestrating all phases of inflammatory response^37, 38^, including producing inflammatory mediators, initiating inflammation, triggering adaptive immunity, resolving inflammation, and restoring homeostasis^39, 40^. Monocytes exhibit high flexibility and diversity in response to environmental cues, allowing them to migrate to inflammatory sites and transform into pro-inflammatory macrophages that release inflammatory cytokines, contributing to both local and systemic inflammation^38, 42^. The abundance of inflammatory monocytes in PBMCs can serve as a biomarker for various inflammatory diseases^41^. When exposed to high-altitude hypoxia, we found the proportions of monocytes and macrophages increased most significantly among these 64 types of immune cells (Fig. 1D and 1E), along with activation of inflammation-related signaling in human monocytic cells (Fig. 1C and 1D). Moreover, the expression levels of inflammatory cytokines (IL-1β, MCP-1, and IL-6) in monocytes-enriched PBMCs significantly increased on the 3-day acute exposure, but gradually faded on 30^th^ prolonged hypoxic exposure of 5500 m (Fig. 1C-1E). In addition, inhibiting the inflammatory response in monocytes significantly decreased the plasma levels of inflammatory cytokines and alleviated the infiltration of inflammatory cells into pulmonary vasculature under acute hypobaric hypoxia (Fig. 7A-7D). These findings suggested that the inflammatory monocytes promptly contribute to the circulating inflammation during hypobaric hypoxia.

### Mitochondrial unfolded protein response induced mito-inflammation in monocytes during hypobaric hypoxia

In response to stress, the permeability of cellular organelles may be altered, allowing certain endogenous molecules known as DAMPs to gain access to PRRs and then initiate inflammatory signaling^14^. The dual membrane structure of mitochondria helps segregate mitochondrial DAMPs from their PRRs, making mitochondrial dysfunction a key player in various inflammatory diseases^15–17^. Under high-altitude hypoxia, we found the mitochondrial protein homeostasis were severely impaired in human circulating leukocytes (Fig. 2A and 2B). A eukaryotic mitochondrion contains approximately 1500 proteins, most of which are encoded by nuclear genes and then translocated from the cytosol to the mitochondrial matrix^43^. To maintain mitochondrial protein homeostasis, molecular chaperones and proteases are cooperated to ensure appropriate folding of newly-imported proteins and degrading of misfolded or non-functional proteins in the mitochondrial matrix^43^. Certain stress conditions, such as the metabolic stress and hypoxia, often disrupt mitochondrial protein homeostasis, leading to the accumulation of unfolded or misfolded proteins in the matrix^44^. Mitochondria are sensitive to the decreased oxygen availability. Therefore, oxygen deprivation, or hypoxia, can impact mitochondrial morphology, mass, and protein compositions, leading to disruptions in mitochondrial metabolism and redox homeostasis, ultimately triggering mitochondrial dysfunction and protein homeostasis^18^. Mitochondrial chaperones HSP60, mitochondrial protease LONP1, and AFG3L2 are three critical effectors of UPR^mt^, which could serve as the checkpoints of mitochondrial stress. From human subjects and mouse model studies, we found both the markers of oxidative stress (SOD, MDA and 8-OHdG) in plasma and the indicators of UPR^mt^ (*Lonp1*, *Afg3l2*, and *Hsp60*) in monocytes-enriched PBMCs initially significantly increased in response to acute hypobaric hypoxia, but gradually decreased to the baseline level in the prolongation of exposure durations (Fig. 2C, 2D and 2F). Additionally, acute hypobaric hypoxia led to noticeable mitochondrial dysfunction characterized by increased mitochondrial ROS production and mitochondrial membrane potential depolarization in circulating monocytes (Fig. 2H). As the initial response to stress, activation of UPR^mt^ is considered to transmit the stress signaling to nucleus and promoted the expression of mitochondrial chaperones and proteases to alleviate mitochondrial proteotoxic stress and restore mitochondrial homeostasis^19, 45^. However, prolonged or continuous activation of UPR^mt^ would increase the expressions of inflammatory genes, enhance cell apoptosis threshold, and maintain heterogeneous mtDNA, ultimately leading to maladaptation^19, 46^.

Inflammasome is a complex inflammatory signaling platform composed of NLRP3, apoptosis-associated speck-like protein containing a caspase activation recruitment domain (ASC), and pro-caspase-1, responsible for the release of IL-1β and IL-18 through caspase-1 activation^47^. Inflammasomes activation is essential for host protection, but dysregulation of inflammasomes would trigger hyperinflammation and pyroptosis^17^. Many mitochondria-dependent molecules have a cross-talk with the inflammasome signaling^17, 44, 48–50^, in which mitochondrial stress of mitochondrial proteotoxicity was reported to directly initiate NLRP3 inflammasome activation in the primary human trophoblasts^51^ and monocytes/macrophages^52^. In our present study, the expression levels of NLRP3 and IL1β significantly increased in PBMCs of human subjects and mouse models when exposed to acute hypobaric hypoxia, but decreased gradually when exposed to prolonged hypobaric hypoxia (Fig. 2E and 2G). In vitro, the signaling of NLRP3 pathways, including NLRP3 expression, and the cleavage of pro-Caspase-1 and pro-IL-1β, were also activated in the monocytes under 1% oxygen hypoxic condition (Figure 3F and 3G, 3J and 3K). These findings indicated the mitochondrial stress, characterized by UPR^mt^ activation and mitochondrial dysfunctions (ROS production and MMP depolarization), could trigger NLRP3 inflammasome signaling and initiate inflammatory response in monocytes under acute hypobaric hypoxia.

### Activation of *Bmal1* drove the mito-inflammation in monocytes by promoting mitochondrial fission during hypobaric hypoxia

In order to achieve optimal fitness, most of organisms anticipate the daily fluctuations in their surroundings, such as light, temperature and food availability. As a result, they exhibit noticeable cyclic fluctuations in behavior and physiology, which is termed as circadian rhythmicity^21^. The molecular clocks provide temporal control and optimize the timing of essential cellular and physiological processes^53^. It is reported that many inflammatory conditions are associated with dysfunctional molecular clocks^54^. Circadian clock also has an important role in controlling mitochondrial metabolism, redox homeostasis and mitochondrial dynamics^25–27^. In the present study, we observed the increased abundance of circulating monocytes were positively correlated with the activation of circadian gene *Bmal1* at high altitude (Fig. 3A). When exposed to hypobaric hypoxia, the fluctuating expression of *Bmal1* in PBMCs were consistent with the dynamic inflammatory response and mitochondrial stress, in which *Bmal1* expression significantly increased under acute hypobaric hypoxia but gradually decreased with prolonged exposure (Fig. 3C and 3D). Additionally, overexpression of *Bmal1* was shown to stimulate and aggregate the hypoxia-induced mitochondrial stress and NLRP3 inflammasome. Conversely, deleting *Bmal1* alleviated the mitochondrial stress and mito-inflammation signaling in monocytes under hypoxia in vivo and in vitro (Fig. 3E-3K, 7B). These results suggested that *i*n response to acute hypobaric hypoxia, upregulation of circadian gene *Bmal1* led to the activation of mitochondrial stress and mito-inflammation in monocytes. Intriguingly, some previous studies reported that downregulation or deletion of *Bmal1* in monocytes and macrophages could increase inflammatory cytokines expression and lead to anti-inflammatory cytokines decreasing^53, 55, 56^, which were not entirely consistent with our present findings. It is well known that the molecular clock keeps time by a series of interlocking positive and negative feedback loops, in which the oscillation of BMAL1: CLOCK heterodimer binding to specific genome sites leads to circadian expression of clock-controlled genes. The expression of *Bmal1* is usually high during the day but repressed during the night^53^. Thus, excessive or insufficient levels of Bmal1 disrupt the normal circadian rhythm and result in adverse consequences.

The remodeling of mitochondrial size and shape, known as “mitochondrial dynamics” including two key processes of fission and fusion^17^. The balance between fission and fusion is crucial for maintaining a healthy population of mitochondria^17^. Mitochondrial fission is a process accounting for mitochondrial fragmentation and generation of smaller mitochondria from larger precursors. This process helps to subdivide mitochondrial population for cell replication and prepares the precondition for dysfunctional mitochondria^17^. It is reported that the dynamic equilibrium of mitochondrial fission–fusion cycle is highly sensitive to hypoxic stress. Under hypoxic conditions, mitochondrial fission is activated to facilitate the selective removal of damaged mitochondria by mitophagy^57, 58^. In cardiomyocytes and hepatocytes, BMAL1 could regulate mitochondrial fission partially by binding to the E-Box elements in the promoter of mitochondrial dynamics genes *Fis1*, *Bnip3*, *Pink1*, and *Mtfr1*^59, 60^. In our present work, we found the mitochondrial fission was enhanced while the mitochondrial fusion was inhibited in monocytes under acute hypobaric hypoxia, along with the impairment of mitochondrial protein homeostasis and activation of *Bmal1* expression (Fig. 2B, 4A and 4B). ChIP assay and *Bmal1* overexpression/knock-out experiments furtherly confirmed that BMAL1 could bind to the E-Box element in the promoter region of *Fis1* gene, and thus enhance mitochondrial fission by promoting *Fis1* transcription in monocytes (Figure 4C-4H). And inhibition of mitochondrial fission by Mdivi-1 could alleviate the BMAL1 overexpression-induced mitochondrial stress and mito-inflammation in monocytes under both normal oxygen and hypoxic conditions (Fig. 5A-5E), suggesting that activation of BMAL1 induced the mitochondrial stress and mito-inflammation by promoting Fis-1 mediated mitochondrial fission in monocytes.

### BHLHE40, a stress-responsive transcription factor, stimulated *Bmal1* transcription in monocytes during hypobaric hypoxia

The circadian clock is continually influenced by environmental cues known as zeitgebers. After decades of studying, light, temperature, food, exercise and mechanosensory stimulation have been identified to the zeitgebers. Disruption of circadian homeostasis results in the negative impacts on human health^21^. Recently, circadian clock has been demonstrated to engage a crosstalk with hypoxic signaling^23^ for the changes in oxygen levels could reset circadian clock. In addition, the core clock component BMAL1 and hypoxia inducible factor (HIF1α) are considered to belong to the same bHLH-PAS TF superfamily^22^. ChIP-sequence analysis has provided insights into the interactions between HIF1A and BMAL1 at a genomic level, revealing a synergistic crosstalk between the two proteins^23^.

Histone modifications and transcriptional factors (TFs) are two crucial mechanisms for controlling gene transcription. The KDM5 family comprises histone demethylases, which are oxygen-dependent dioxygenases that regulate gene transcription epigenetically by targeting H3K4me1/2/3. H3K4me1/2/3 was predicted to occur in the regulatory region of *Bmal1* gene and we observed the H34Kme3 level was significantly increased in monocytes under hypoxia both in vivo and in vitro (Fig. 6B and 6C). Interestingly, inhibiting the activity of KDM5 demethylases or reducing the expression of KDM5A/5C genes did not enhance Bmal1 transcription (Fig. 6D and 6E). Based on TF prediction (Table 2), hypoxic DEGs (Fig. 6F and 6G), and IPA analysis (Table 3), the transcriptional factor BHLHE40 was screened to have the potential regulation in *Bmal1* transcription under hypoxia (Fig. 6H). Furthermore, we observed a significant increase in the mRNA expression of Bhlhe40 under hypoxic conditions in vivo and in vitro (Fig. 6J and K). Inhibition of Bhlhe40 led to an obvious decrease in Bmal1 transcription (Fig. 6I) and alleviated the mitochondrial stress and inflammatory response under hypoxia (Fig. 6I, 6L, and 6M). As a member of the basic helix-loop-helix TF family, BHLHE40 was reported to bind DNA at class B E-box motifs and functions^61^. BHLHE40 can be induced in many stress conditions, such as hypoxia and ER stress, and is involved in regulating cell cycling, cell death, differentiation, and cytokine production ^61, 62, 66^. BHLHE40 is an important hypoxia-induced tumors marker and its expression is directly regulated by HIF-1α^63–65^ (Fig. S3B). Additionally, BHLHE40 could act a negative feedback by repressing the CLOCK: BMAL1 transactivation with the direct protein-protein interactions or competition for the E-box element^67, 68^. Under hypobaric hypoxia, we found BHLHE40 served as an upstream regulator of *Bmal1* transcription, which drove mitochondrial stress and inflammatory response by enhancing *Bmal1* expression in monocytes.

### Knock-out of myeloid-specific *Bmal1* alleviated the inflammatory activity of PBMCs during acute hypobaric hypoxic stress

The molecular clocks are present in almost all cells of the body. Dysfunctional molecular clocks in immune cells have been linked to inflammatory diseases^56^. In myeloid cells, BMAL1 has been demonstrated to impact monocyte diurnal oscillations^28^, macrophage immune functions^69^, and inflammatory cytokine production^28, 56, 70^. Absence of an intrinsic clock may have beneficial effects under certain conditions^71^. For example, *Bmal1* deletion in vascular smooth muscle cells protected mice from abdominal aortic aneurysm^72^. In addition, *Bmal1* deletion in myeloid cells reduced aortic inflammatory response and retarded atherogenesis^71^. Global deletion of *Bmal1* in adult mice has been shown to facilitate the adaptation to light/dark disruption and then protect mice from the insulin resistance and atherosclerosis^73, 74^. BMAL1 is essential for normal circadian rhythm in physiology and behavior, however, lacking intrinsic clock may render organism less vulnerable to environmental disturbances and then have the positive outcomes^75^. We found that myeloid-specific *Bmal1* knout-out alleviated the systemic circulating inflammation and vascular inflammation under acute hypobaric hypoxia (Fig. 7 and Video S1-S3), which suggested that *Bmal1* deletion may exert beneficial effects in certain extreme environments.

### Practical implications and limitations

Hypoxia-induced inflammation has been considered to plays an important role in the development of high-altitude illnesses, however, the origin of inflammatory cytokines, the specific responding cell types, and the signaling mechanisms remain unclear. In addition, the dynamic changes of inflammatory response at different altitudes and different exposure durations are rarely reported. From human subjects and mouse models, our present work indicated a dynamic inflammatory response in circulating plasma and PBMCs at 3-day and 30-day exposure of 5500 m hypobaric hypoxia, in which inflammatory monocytes promptly contribute to the circulating inflammation. Furthermore, our present work raised a novel mechanism by which mitochondria transmit the hypoxic signaling by activation of circadian gene Bmal1and then control the inflammation. Therefore, it is thought that therapeutic manipulations targeting circadian rhythm, and mitochondrial stress in monocytes may represent an opportunity to treat clinical hypoxia-related diseases. There is a limitation in our present study. The inflammation in monocytes faded gradually with the prolongation of hypobaric hypoxic exposure, but the inflammatory cytokines in plasma remained continuous upregulation. We supposed that in addition to immune cells, there may be other cells like endothelial cells and vascular smooth muscle cells, continuously contributing to the circulating inflammation in response to hypobaric hypoxia.

## Conclusions

In summary, BHLHE40, a transcription factor associated with hypoxia, stimulated *Bmal1*, which in turn triggered the mitochondrial unfolded protein response and drove the mito-inflammation in monocytes by promoting Fis1-mediated mitochondrial fission. Our work provides a novel mechanism which may develop the circadian targeting drugs for altitude or hypoxia-related diseases.

## Acknowledgements

M-J.X., Y-Z.Y., Y-L.G. and Y.L. designed and performed the experiments, prepared the figures, and wrote the manuscript. B.Z., J.X., S-Y.H., Q-L.C., and P-J.L. contributed to the performance of the experiments. Y-R.B., Y-G.B., and L.Z. supervised the work and wrote the manuscript.

## Sources of Funding

This study was supported by grants from the National Natural Science Foundation of China (82072101) and the National Key Research and Development Program of China (2020YFA0803603 and 2021YFA1301402).

## Supplemental information

Supplemental figures: Figure S1-S3

Supplemental videos: Video S1-S3

Supplemental tables: Table S1-S3

## Novelty and Significance

### What Is Known?

Acute mountain sickness (AMS) is a common risk for sojourners and mountaineers at high altitude, which can progress to severe and potentially fatal illnesses in some cases. Hypoxic stress-induced inflammation is implicated in the onset and progression of AMS. However, the origin of inflammatory cytokines, the specific responding cell types, and molecular mechanisms remain unknown.
λ Mitochondria are essential organelles of cellular functions and metabolism. In addition, Mitochondrial could transmit the signaling pathways to control the inflammation, which is known as a new concept of the mito-inflammation.
λ Mitochondrial functions are under circadian regulation and hypoxia significantly disrupts circadian rhythm during environmental adaptation.

### What New Information Does This Article Contribute?

λ Monocytes in peripheral blood mononuclear cell (PBMCs) were more sensitive and contributed promptly to circulating inflammation in response to acute hypobaric hypoxia.
λ Hypoxic stress triggered the mitochondrial unfolded protein response and then induced the mito-inflammation (NLRP3 inflammasome) in monocytes.
λ Activation of Bmal1 drove mitochondrial stress and mito-inflammation in monocytes by promoting Fis1-mediated mitochondrial fission.
λ BHLHE40, a stress-responsive transcription factor directly targeted by HIF-1α, stimulated Bmal1 transcription in monocytes under hypobaric hypoxia.
λ Myeloid-specific Bmal1 deletion alleviated systemic circulating and vascular inflammation under acute hypobaric hypoxia.

## Notes

### Competing Interest Statement

The authors have declared no competing interest.

